# Structural insights into terminal arabinosylation biosynthesis of the mycobacterial cell wall arabinan

**DOI:** 10.1101/2024.09.17.613533

**Authors:** Yaqi Liu, Chelsea M. Brown, Satchal Erramilli, Yi-Chia Su, Po-Sen Tseng, Yu-Jen Wang, Nam Ha Duong, Piotr Tokarz, Brian Kloss, Cheng-Ruei Han, Hung-Yu Chen, José Rodrigues, Margarida Archer, Todd L. Lowary, Anthony A. Kossiakoff, Phillip J. Stansfeld, Rie Nygaard, Filippo Mancia

## Abstract

The emergence of drug-resistant strains exacerbates the global challenge of tuberculosis caused by *Mycobacterium tuberculosis* (*Mtb*). Central to the pathogenicity of *Mtb* is its complex cell envelope, which serves as a barrier against both immune system and pharmacological attacks. Two key components of this envelope, arabinogalactan (AG) and lipoarabinomannan (LAM) are complex polysaccharides that contain integral arabinan domains important for cell wall structural and functional integrity. The arabinofuranosyltransferase AftB terminates the synthesis of these arabinan domains by catalyzing the addition of β-(1→2)-linked terminal arabinofuranose residues. Here, we present the cryo-EM structures of *Mycobacterium chubuense* AftB in its apo and donor substrate analog-bound form, determined to 2.9 Å and 3.4 Å resolution, respectively. Our structures reveal that AftB has a GT-C fold transmembrane (TM) domain comprised of eleven TM helices and a periplasmic cap domain. AftB has an irregular tube-shaped cavity that bridges the two proposed substrate binding sites. By integrating structural analysis, biochemical assays, and molecular dynamics simulations, we elucidate the molecular basis of the reaction mechanism of AftB and propose a model for catalysis.

## INTRODUCTION

Tuberculosis (TB), caused by *Mycobacterium tuberculosis* (*Mtb*), is the world’s second deadliest infectious disease, surpassed only by COVID-19 during the recent pandemic^1^. The resilience and virulence of *Mtb* is largely attributed to its unique cell envelope, a lipid-rich fortress that confers resistance to host immune mechanisms and provides an impenetrable barrier to a spectrum of antibiotics^2–5^. This cell envelope is comprised of three major components: long-chain mycolic acids creating a protective lipid layer, peptidoglycan providing cell shape and rigidity, and an arabinogalactan (AG) domain linking the two (Fig. 1a). AG is a complex polysaccharide characterized by a linear galactan backbone and a highly branched arabinan domain (Fig. 1a). Together, these three components form the mycolyl–arabinogalactan–peptidoglycan (mAGP) complex, which provides an impermeable shield crucial for survival and pathogenicity^6–9^ (Fig. 1a). In addition to the mAGP complex, the cell envelope contains a myriad of glycolipids, such as phosphatidylinositol mannosides (PIMs) and their hyperglycosylated derivatives lipomannan (LM) and lipoarabinomannan (LAM) as well as a delipidated form – arabinomannan (AM). These species are important for growth, virulence, and interaction with the host immune system^9,10^. Among these, LAM/AM possesses a complex structure, containing a mannan core and a highly branched arabinan domain^10–14^ (Extended Data Fig. 1a).

**Figure 1.**
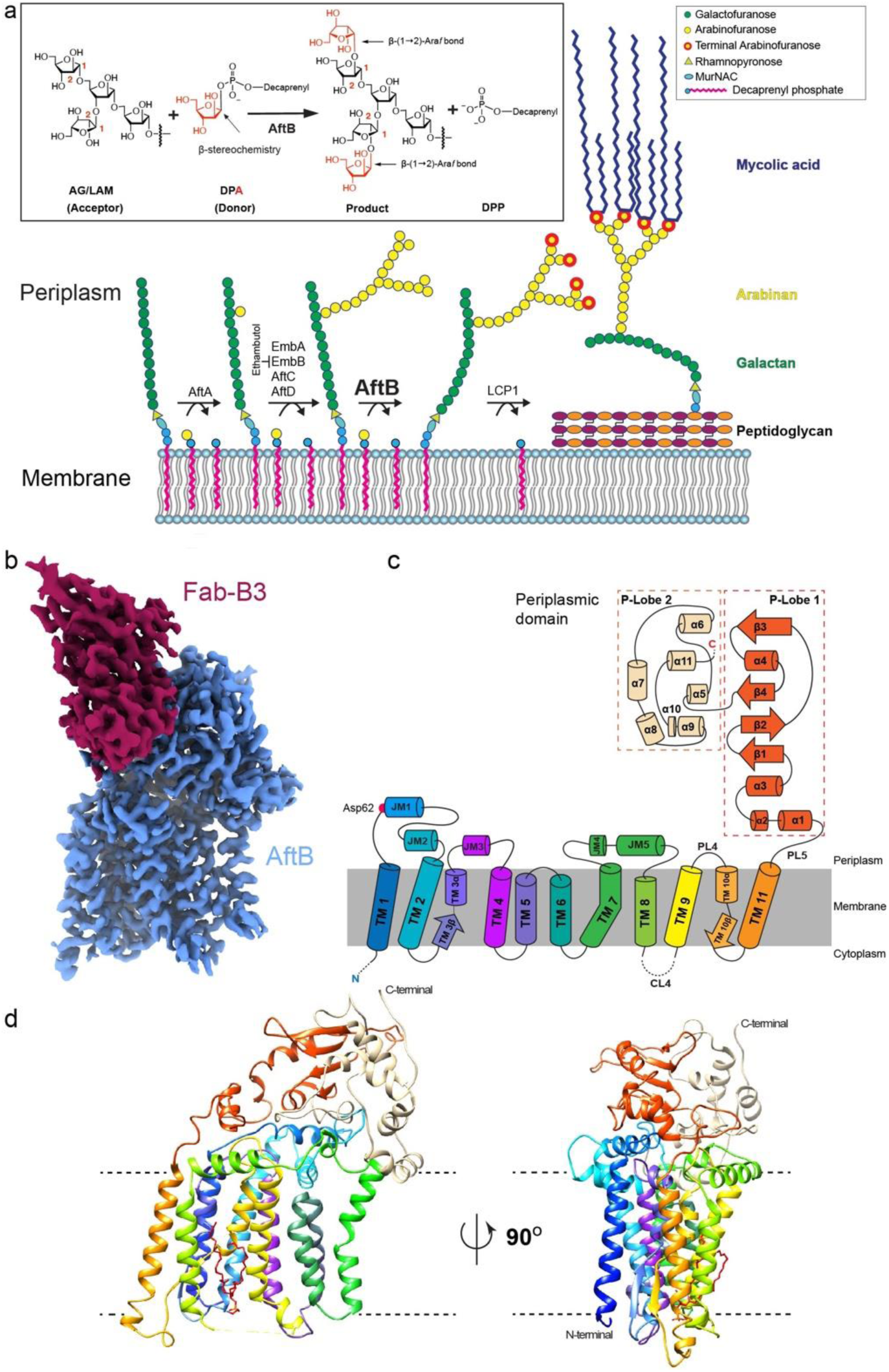
Biosynthetic Pathway and Structural Architecture of AftB. (a) Schematic of the arabinogalactan biosynthetic pathway in mycobacteria, featuring the enzymatic reactions carried out by AraTs. Enzyme names are indicated above the reaction arrows. The inset shows the β-(1→2) arabinosyl transfer reaction catalyzed by AftB, (b) Cryo-EM density map of AftB in complex with Fab-B3. AftB is shown in blue, while Fab-B3 is shown in maroon. (c) Topological diagram of AftB showing the arrangement of secondary elements in the TM region and in the PD, annotated and colored in correspondence with Fig 1(d). The catalytic residue Asp62 is denoted as a red dot. (d) Structure of AftB shown in ribbon. The unmodeled region of AftB is marked as dashed lines, and membrane boundaries are marked by dotted horizontal bars.

The distinctive characteristics of the mycobacterial cell envelope have established it as a widely exploited target for anti-TB therapeutics^5,6,15–17^. Front-line anti-TB drugs such as isoniazid and ethambutol specifically target *Mtb* cell envelope biosynthesis^18,19^. Isoniazid disrupts the synthesis of mycolic acids, a component of the mAGP^18–20^ complex, whereas ethambutol targets enzymes involved in arabinan biosynthesis for AG and LAM^21–24^. However, the emergence of drug-resistant and multidrug-resistant strains of *Mtb* has underscored the limitations of current TB treatments and highlighted the pressing need for innovative therapeutic drug targets and strategies^5,25,26^.

The biosynthesis of the arabinan domains of AG and lipoarabinomannan LAM involves a series of reactions catalyzed by arabinofuranosyltransferases (AraTs)^5,6^. These membrane-embedded glycosyltransferases orchestrate the addition of arabinofuranose (Ara*f*) from β-D-arabinofuranosyl-1-monophosphoryldecaprenol (DPA), the only proven D-arabinose donor, onto the growing polysaccharide chains^27–29^. For AG, the biosynthesis of the arabinan domain begins with the α-(1→5)-arabinosyltransferase AftA, which adds the first α-D-Ara*f* residue to the mature galactan backbone^30^ (Fig. 1a). This is followed by the action of enzymes EmbA and EmbB, which extend the arabinan chain through α-(1→5) linkages^31^ (Fig. 1a). For LAM, the priming enzyme responsible for initiating arabinan synthesis remains unidentified. Nonetheless, it is known that EmbC is involved in the extension of arabinan-primed LM with α-(1→5)-Ara*f* residues^32^. AftC introduces α-(1→3)-branching within both AG and LAM, adding structural complexity to these components of the cell envelope^33^(Fig. 1a). AftD is thought to have a similar function to AftC, yet its precise role remains not fully elucidated^34,35^. Finally, AftB, an integral membrane enzyme encoded by the gene Rv3805c in *Mtb*, catalyzes the addition of the terminal β-(1→2)-linked D-Ara*f* residues to both AG and LAM^36,37^ (Fig. 1a and Extended Data Fig. 1a). In AG, this results in a terminal hexa-arabinofuranoside motif (Ara*f*_6_) at the nonreducing end, which provides the anchoring point for mycolic acid^6,38^ (Fig. 1a). A ligating phosphotransferase termed Lcp1 attaches AG to the peptidoglycan, completing the AG assembly^39^ (Fig. 1a).

AftB is thought to play an important role in maintaining the integrity and functionality of the mycobacterial cell envelope and has emerged as a promising drug target for TB treatment^36,40,41^. However, the structural basis of the molecular mechanisms of AftB have yet to be fully elucidated and, to date, there are no published structures of this enzyme. Here, we present structures of *Mycobacterium chubuense* AftB in both its apo and substrate analog-bound forms, determined by single-particle cryogenic-electron microscopy (cryo-EM). Using a combination of this structural information, biochemical assays, and molecular dynamics (MD) simulations, we identified the substrate binding sites and proposed a mechanism of action for the enzyme.

## RESULTS

### Structure determination of AftB

We screened AftB orthologs from 45 mycobacterial species for expression in *E. coli* and identified AftB from *Mycobacteria chubuense* (*Mc*AftB) as the most promising target for structural determination. To confirm that recombinant AftB is active, we adapted a reported arabinosyltransferase assay^42^ using a tetrasaccharide acceptor substrate (Extended Data Fig. 2a) corresponding to the non-reducing terminus of the natural AftB substrate in AG and LAM, and farnesyl phosphoarabinose (FPA) as surrogates for natural substrates (Extended Data Fig. 2b). FPA has been shown previously to be an effective donor substrate for mycobacterial AraTs, and is much simpler to prepare and handle than DPA^42,43^. Mass spectrometry analysis of the acetylated products confirmed that AftB catalyzes the transformation of the tetrasaccharide first to pentasaccharides and ultimately to a hexasaccharide, thus confirming enzymatic activity (Extended Data Fig. 2c-d).

The 78 kDa *Mc*AftB, was purified in detergent and reconstituted into lipid-filled nanodiscs for structure determination by cryo-EM (Extended Data Fig. 1b-c). To provide fiducials for particle alignment and increase particle size to facilitate structure determination, we screened a synthetic phage display library to select recombinant antigen-binding fragments (Fabs) against *Mc*AftB^44,45^. Seven high-affinity Fab candidates were identified and evaluated for their ability to form complexes with *Mc*AftB, and Fab-B3 was selected due to its high binding affinity (Extended Data Fig. 1d-e).

We collected 7,164 micrographs of nanodisc-reconstituted apo *Mc*AftB in complex with Fab-B3. After iterative 2D classification, we obtained high-quality 2D class averages showing clear features for the transmembrane (TM) domain and bound Fab. After further sorting the particles using three-class *ab initio* modeling, we obtained a map with an overall resolution of 2.9 Å (Extended Data Fig. 3). This allowed us to build an almost complete atomic model of *Mc*AftB, apart from 28 residues at the N-terminus, 11 at the C-terminus, and a disordered region (323–334) in a cytoplasmic loop connecting two TM helices (8 and 9) (Fig. 1 and Extended Data Fig. 4). The variable region of the Fab fragment was well resolved, and we could build it reliably, providing a clear depiction of the interaction interface (Fig. 1 and Extended Data Fig. 1f-g). On the other hand, the constant domain of the Fab was omitted from the model due to the disordered nature of the corresponding region in our density map.

Additionally, we observed a distinct diacyl-glycerophospholipid-like density between TM helices 8 and 9. We tentatively attributed this to phosphatidylethanolamine (PE), given its abundance in *E. coli* membranes^46^ (Extended Data Fig. 4).

### Overall structure of AftB

AftB contains two distinct domains: a TM domain with 11 α-helices and a C-terminal periplasmic domain (PD) composed of a mixture of α-helices and β-strands (Fig. 1c-d). The TM helices of AftB are connected by five cytoplasmic loops, CL1–CL5, and four periplasmic loops, PL1–PL4. All cytoplasmic loops are well-resolved in the structure, except for CL4 (residues 323–334), which connects TM helices 8 and 9 (Fig. 1c-d). Three of the four periplasmic loops have a more complex structure. PL1, located between TM helices 1 and 2, features two juxtamembrane (JM) helices, JM helix 1 and 2. PL2, which connects TM helices 3 and 4, contains JM helix 3, while PL3, spanning between TM helices 7 and 8, harbors JM helices 4 and (Fig. 1c-d). The C-terminal domain of AftB, extending from the end of the eleventh TM helix, forms a dome-like structure in the periplasmic space, capping the TM helical bundle (Fig. 1b, 1d). This domain is composed of two lobes: the first consists of four β-strands (β1–β4) and four α-helices (α1–α4), along with connecting loops, and the second lobe contains seven α-helices (α5–α11), with loops between them. PL5 connects TM11 to α1, serving as a linker between the TM bundles and the PD (Fig. 1d).

Fab-B3 binds to the PD of AftB, with both its light chain and heavy chain contributing to the interaction interface (Extended Data Fig. 1f). The AftB–Fab-B3 interface is dominated by a set of hydrogen bond interactions mediated primarily by the heavy chain of the antibody fragment (Extended Data Fig. 1g).

### Conserved GT-C fold of AftB

Glycosyltransferases are structurally diverse enzymes that can be classified into three main fold types: GT-A, GT-B, and GT-C. While GT-A and GT-B folds typically contain Rossmann-like domains, GT-C fold enzymes are characterized by multiple TM helices and can be further classified into GT-C_A_ and GT-C_B_ subclasses^47,48^. Using the DALI server^49^, we found that AftB belongs to the GT-C_A_ superfamily of glycosyltransferases^47^. The first seven helices of the TM domain have a GT-C glycosyltransferase fold^47,50^. The enzymes with the highest structural homology to AftB include human oligosaccharyltransferase complex catalytic subunit B (STT3B)^51^, bacterial oligosaccharyltransferase PglB^52,53^, bacterial aminoarabinose-transferase ArnT^53^, and yeast glucosyltransferase ALG6^54^ (Extended Data Fig. 6). These all belong to the GT-C_A_ class^47^ and share a conserved N-terminal module with AftB consisting of the first seven TM helices 1 and 7, including two JM helices (JM1 and JM2) between TM helices 1 and 2, followed by the structurally variable additional TM helices. While adopting the overall GT-C_A_ fold in its N-terminal module, AftB exhibits notable differences within this core fold. In AftB, both TM helices 3 and 10 are characterized by a split into a short α-helix and a β-strand, with these β-strands interacting through hydrogen bonds to form a small anti-parallel β-sheet within the TM domain (Fig. 1c-d and Extended Data Fig. 4b). In contrast, other GT-C_A_ enzymes typically display a kink in TM3, splitting it into two helices. This unique structural feature may potentially provide enhanced structural flexibility, contributing to AftB’s specific substrate recognition or catalytic mechanism.

### The putative substrate cavity of AftB

We observed an irregular tube-shaped cavity between the TM domain and the PD, with one end facing the membrane and the other facing the periplasmic space (Fig. 2a-b). Notably, all GT-C_A_ enzymes with high structural homology to AftB contain a catalytically essential aspartate located at the tip of JM1^51–54^. Asp62, which belongs to a conserved DD motif, is located at the tip of JM helix 1 and is buried inside this cavity (Fig. 2a), suggesting its potential role as the catalytic residue. To further investigate the importance of Asp62, we generated a D62A mutant and assessed its enzymatic activity. We confirmed comparable expression levels of the D62A mutant with wild-type *Mc*AftB (data not shown), and our arabinosyltransferase assay revealed that it was catalytically inactive, confirming the critical role of Asp62 in AftB function (Extended Data Fig.2e).

**Figure 2.**
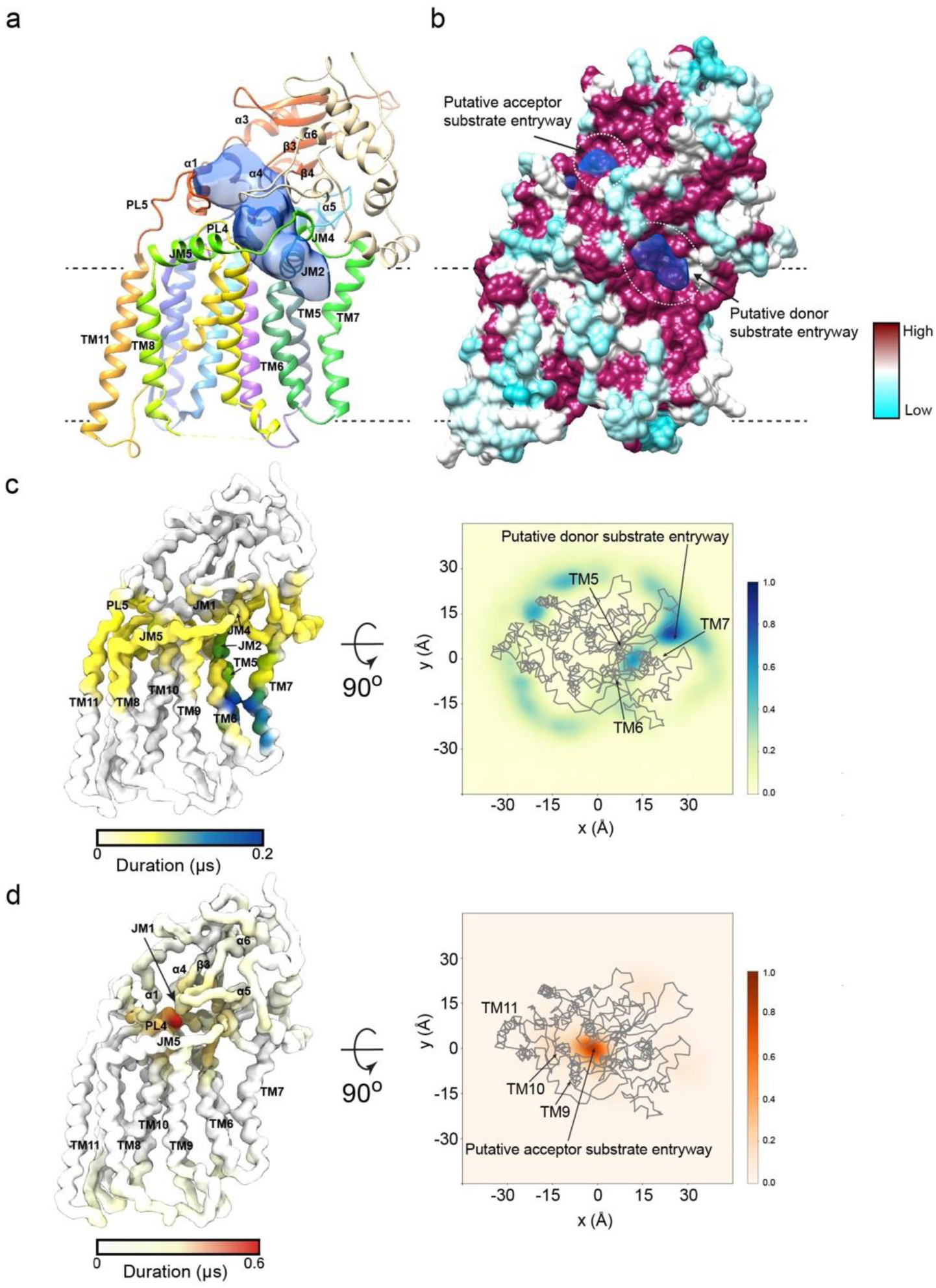
Putative substrate binding cavity. (a) AftB structure shown as ribbon, and colored as in Fig. 1c, with the cavity shown in semi-transparent blue. Volumes were calculated using the Voss Volume Voxelator server^104^ with probe sizes of 2 Å and 8 Å. (b) Surface of AftB by conservation, with a gradient from magenta (absolutely conserved) to cyan (least conserved), cavity shown in semi-transparent blue. Predicted entryways for substrates are marked with dotted circles. (c) Left shows the AftB backbone as a surface, colored by the duration of interaction with DPA in CG-MD simulations. The darker the color, the longer the duration of interaction. The right plot shows the density in the x and y dimensions of DPA relative to the protein shown in gray, where the darker regions show higher density. (d) Shows the same as (c) but for terminal-Ara*f*_4_.

JM helices 1, 2, and 4 and PL4 line the segment of the cavity facing the membrane, alongside the connecting loops between JM helices 1 and 2, TM helices 3 and 4, and JM helices 4 and 5. The cavity extends towards the membrane, with an opening defined by TM helices 5, 6, and 7 (Fig. 2a and Extended Data Fig. 9a and c). The opposite end of the cavity faces the PD and is defined by the loops between TM helix 11 and α1, β3 and α4, β4 and α5, and between α5 and α6. The architecture of the cavity suggests an obvious entry path for the donor substrate DPA, through the opening towards the lipid bilayer. The polar head of DPA would be positioned towards the catalytic core of the enzyme, allowing the lipid tail to extend out into the lipid bilayer. The wider opening of the cavity towards the periplasm presents a predominantly hydrophilic environment, which could accommodate the branched tetrasaccharide domain of the acceptor substrate (Fig. 2a, Extended Data Fig. 8f).

To further substantiate our hypothesis for entry of the acceptor and donor substrates, we performed coarse-grained molecular dynamics (CG-MD) simulations to identify preferential binding sites. We used the native donor substrate DPA (Extended Data Fig. 8a) and a hexasaccharide fragment of the native arabinan acceptor substrate which we term “terminal-Ara*f*_4_”. This acceptor species includes the branched motif present in the acceptor substrate used in the functional assay with two additional α-(1→5)-Ara*f* residues at the non-reducing end (Fig, 1a, Extended Data Fig. 8b). Analyses of our simulations reveal a preferential binding site for DPA within the membrane domain along TM helices 5, 6, and 7, as well as within the region of the cavity framed by JM helices 1, 2 and 4 (Fig. 2c). This binding site overlaps with the cavity observed in the structure and its opening towards the membrane (Fig. 2a-b and Supplementary Fig. 10c). In the CG-MD simulations, the terminal-Ara*f*_4_ showed rapid association from the periplasmic side of AftB from bulk solvent, mostly in a cavity framed by the loops connecting β3 and α4, β4 and α5, and α5 and α6 (Fig. 2d). This corresponds to the predicted entryway for the acceptor substrate (Fig. 2a-b). For comparison, RoseTTAFold All-Atom^55^ was used to fold and dock both substrates and products to AftB, further predicting and characterizing their binding sites.

### The Cryo-EM structure of 2F-FPA-bound AftB

Next, we sought to further investigate the molecular basis of substrate recognition and catalysis by determining the structure of AftB in a substrate-bound form. To this end, we used 2-F-farnesyl phosphoarabinose (2F-FPA), a donor analog for mycobacterial AraT^56^. In 2F-FPA, the decaprenyl chain is replaced with a shorter farnesyl group, making the substrate less hydrophobic than DPA and better suited for structural studies (Extended Data Fig. 2). The incorporation of the fluorine atom into the Ara*f* ring stabilizes the labile glycosyl-phosphate bond in turn making it more amenable as a probe.

We determined the structure of nanodisc-reconstituted AftB in complex with 2F-FPA and bound to Fab-B3 by cryo-EM to an overall resolution of 3.4 Å (Fig. 3, Extended Data Fig. 3d). We were able to build the entire protein except for the 28 residues at the N-terminus, 11 residues at the C-terminus, a disordered region (319–335) at the cytoplasmic loop that connects TM helices 8 and 9, and a small region 432–437 on the periplasmic loop between TM helix 11 and periplasmic helix α1 (Extended Data Fig. 5). The Fab-B3 binds AftB in a similar conformation to that observed in the apo structure (Extended Data Fig. 3d). The density for 2F-FPA is well resolved, showing that this ligand adopts a partially curved conformation with the sugar ring projecting deep into the cavity (Fig. 3a). The hydrophobic acyl chain of 2F-FPA extends along TM helix 6 and the phosphate moiety is coordinated by Arg221 and Arg372 and forms a hydrogen bond with Thr91 located in the loop between JM helices 1 and 2 (Fig. 3b). Thr274 coordinates the fluorine on the arabinofuranose ring, and Glu172 forms a hydrogen bond with the C-5 hydroxyl group of the arabinofuranose (Fig. 3b). Not surprisingly, these interacting residues are conserved across mycobacteria (Extended Data Fig. 7).

**Figure 3.**
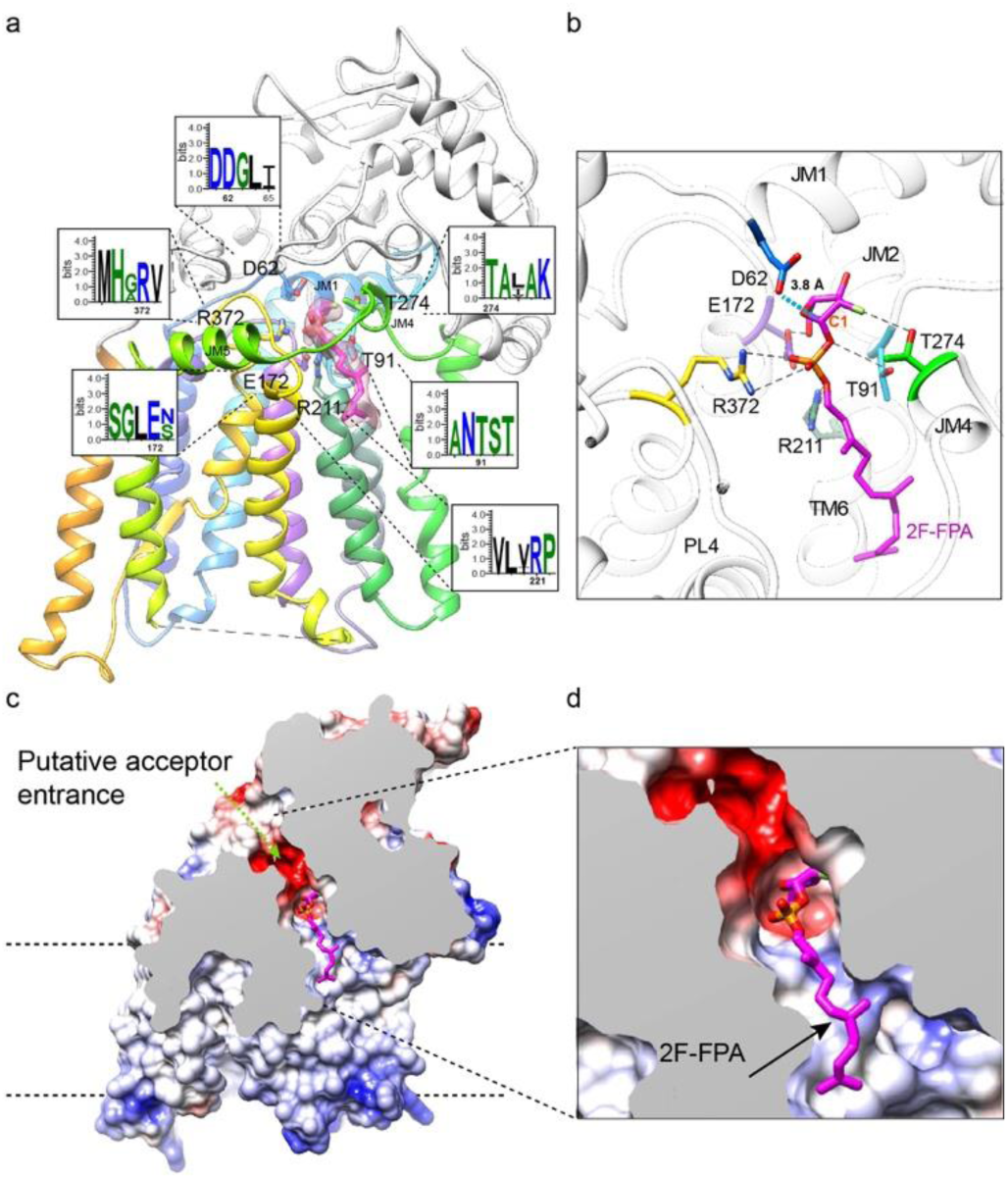
Interaction between AftB and bound 2F-FPA. (a) Structure of AftB complexed with 2F-FPA shown in ribbon. The TM domain of AftB is colored in rainbow as in Fig 1d, while the PD is colored in white. 2F-FPA is shown in sticks and colored in magenta. Cryo-EM density of 2F-FPA colored with semi-semitransparent magenta. WebLogo plots^105^ show the amino acids interacting with 2F-FPA. (b) Close-up view of the interaction between AftB and 2F-FPA. Polar contacts are indicated as black dashed lines. Residues involved in the interaction are shown as sticks. The distance between the catalytic residue D62 and C1 of 2F-FPA is indicated with blue dashed lines. (c) and (d) Electrostatic surface depiction of the AftB-2F-FPA complex, a partial cross-sectional perspective is applied to highlight the active site cavity with the bound donor substrate-analog 2F-FPA.

### Structural Rearrangements upon Substrate Binding

Comparison of the apo and 2F-FPA-bound AftB structures shows conformational changes primarily within the PD. Within the TM domain, TM helices 7 and 11 undergo an inward shift, particularly pronounced in TM11, accompanied by a noticeable shift in loop PL5 (Fig. 4). The shift in PL5 appears to initiate a conformational change in the PD, resulting in a clamp-like movement of it, towards the membrane (Fig. 4).

**Figure 4.**
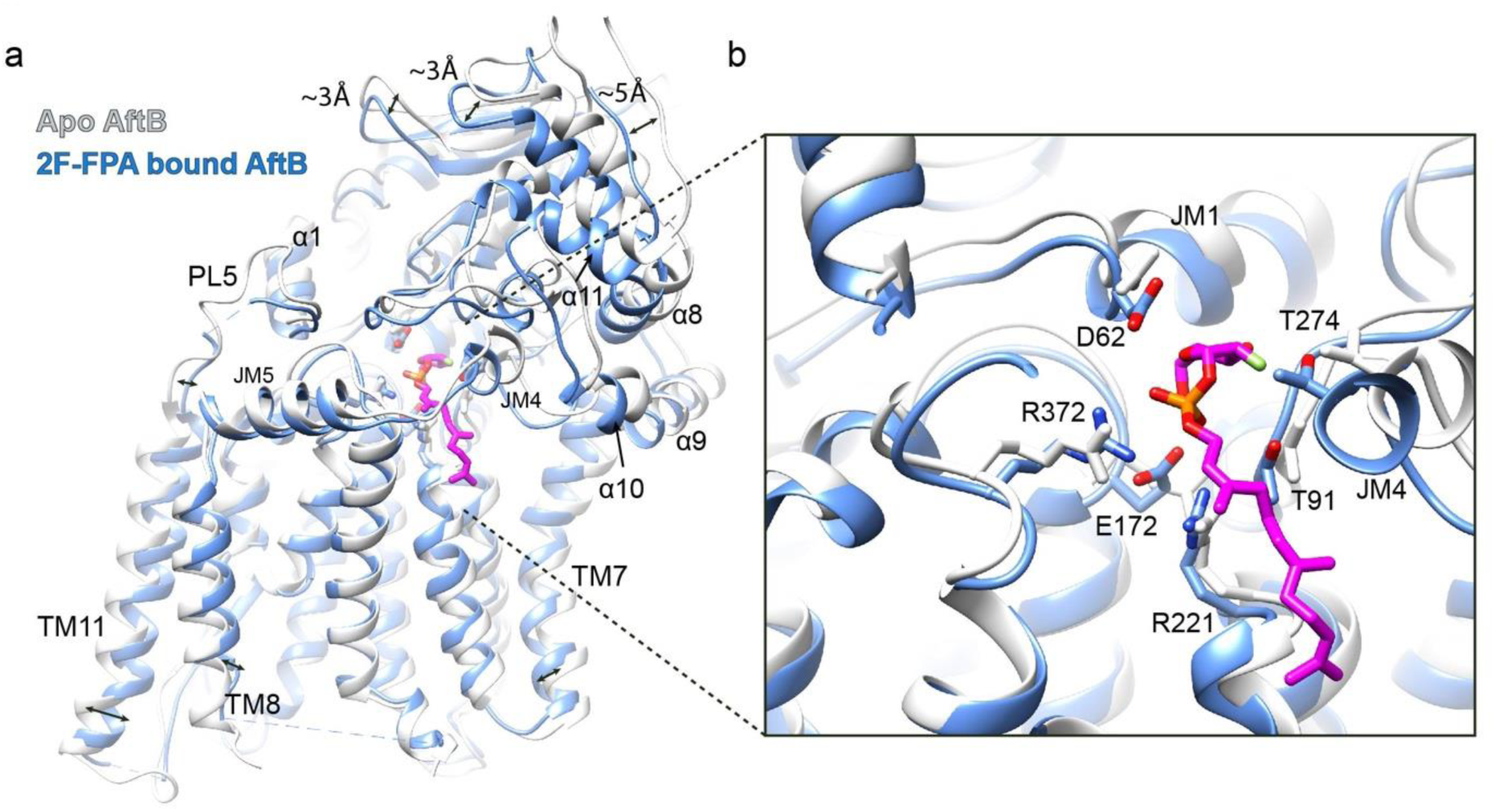
Conformational rearrangement of AftB upon 2F-FPA binding. (a) Superimposition of apo (white) and 2F-FPA bound AftB (blue). PD undergoes a clamp-like movement while TM helices exhibit minor inward movement. (b). Magnified view of the substrate pocket in the apo structure (white) and the 2F-FPA-bound structure (blue), superimposed, showing interacting residues shifted toward 2F-FPA.

A handful of residues within the substrate-binding cavity, in particular Asp62, Glu172, Arg372, Arg221, Thr91, and Thr274, shift towards the bound 2F-FPA, suggesting a coordinated adaptation to facilitate interaction (Fig. 4). The movement of these residues contributes to the slight inward pivoting of TM helices 1 and 7. The interplay between the conformational changes in the TM and PD results in a relatively more compact conformation of AftB.

### Insights into DPA and Acceptor Binding Mode with Docking and MD simulation

To further our understanding of the interaction between the enzyme and its substrates, we used a combination of docking and MD simulations. Ligand docking and RoseTTAFold analysis of the donor substrate propose that DPA binds similarly to the 2F-FPA in the AftB-2F-FPA complex, with its polar head oriented towards Asp62 in the active site (Extended Data Fig. 8d-e). The phosphate group of DPA is coordinated by Arg372 and Arg221, along with interactions involving Thr91 and Thr243, mirroring the coordination seen with 2F-FPA (Extended Data Fig. 8d). At this position, it is coordinated with conserved residues including Phe368 from PL4, and Phe448, Tyr449 from the first periplasmic alpha helix α1, Asp541 from the loop between the third periplasmic beta strand β3 and α4, and Arg560 and His563 from the loops linking β4 to α5 and between alpha helices α5 and α6 (Extended Data Fig. 8e).

To further explore the interaction of the substrates and products of this reaction with AftB, atomistic resolution MD simulations were performed. In these simulations, we used the native donor substrate DPA, its by-product decaprenyl phosphate DP, and two polysaccharide mimics: a terminal-Ara*f*_4_, as described above, representing the acceptor substrate, and a “terminal-Ara*f*_6_” representing the final product (Extended Data Fig. 8a and c). The terminal-Ara*f*_6_ represents the arabinosylated product with the characteristic Ara*f*_6_ motif at the end of the arabinan domain, including two additional α-(1→5)-Ara*f* residues from the linear backbone. We conducted simulations with either the substrates (DPA and terminal-Ara*f*_4_) or the products (DP and terminal-Ara*f*_6_) present. In both simulation set-ups, the terminal-Ara*f*_4_ and terminal-Ara*f*_6_ escaped their bound pose (Extended Data Fig. 9a). We anticipate that these terminal mimics leave the binding site as they lack the lengthy repeating sugar units and the lipid membrane anchors. Conversely, both the lipid donor and its by-product are stable within the binding pocket (Extended Data Fig. 9a) and retain their docked pose (Extended Data Fig. 9b-d). The addition of either substrates (DPA with terminal-Ara*f*_4_) or products (DP and terminal-Ara*f*_6_) does not have any significant effect on the dynamics of the protein when compared to simulations with the apo state (Extended Data Fig. 10a-b). Nevertheless, the p*K*_a_ of the proposed catalytic residue Asp62 lowers substantially when substrates or products are included in the simulation (Extended Data Fig. 10c). This is not observed with other substrate-interacting residues (Extended Data Fig. 10c), which further supports the relevance of Asp62 in enzymatic activity. Based on this, we propose that at physiological pH, Asp62 would predominantly be in its deprotonated, catalytically active form (see below).

### Catalytic Mechanism of AftB

AftB catalyzes the transfer of D-Ara*f* from the donor DPA to the acceptor, creating a β-(1→2) glycosidic bond^36,37,57^ (Fig. 1a). This enzymatic process preserves the β-configuration of the anomeric carbon from the donor to the product, hallmarking AftB as a retaining glycosyltransferase^36,37,57,58^. Our structure reveals that Asp62, located within the conserved DD motif, is positioned approximately 3.8 Å away from the anomeric carbon (C1) of 2F-FPA, further strengthening its suggested role as a catalytic nucleophile in the reaction. Our structural and computational studies provide a model for how AftB brings together the donor DPA and acceptor substrate for catalysis (Fig. 5a).

**Figure 5.**
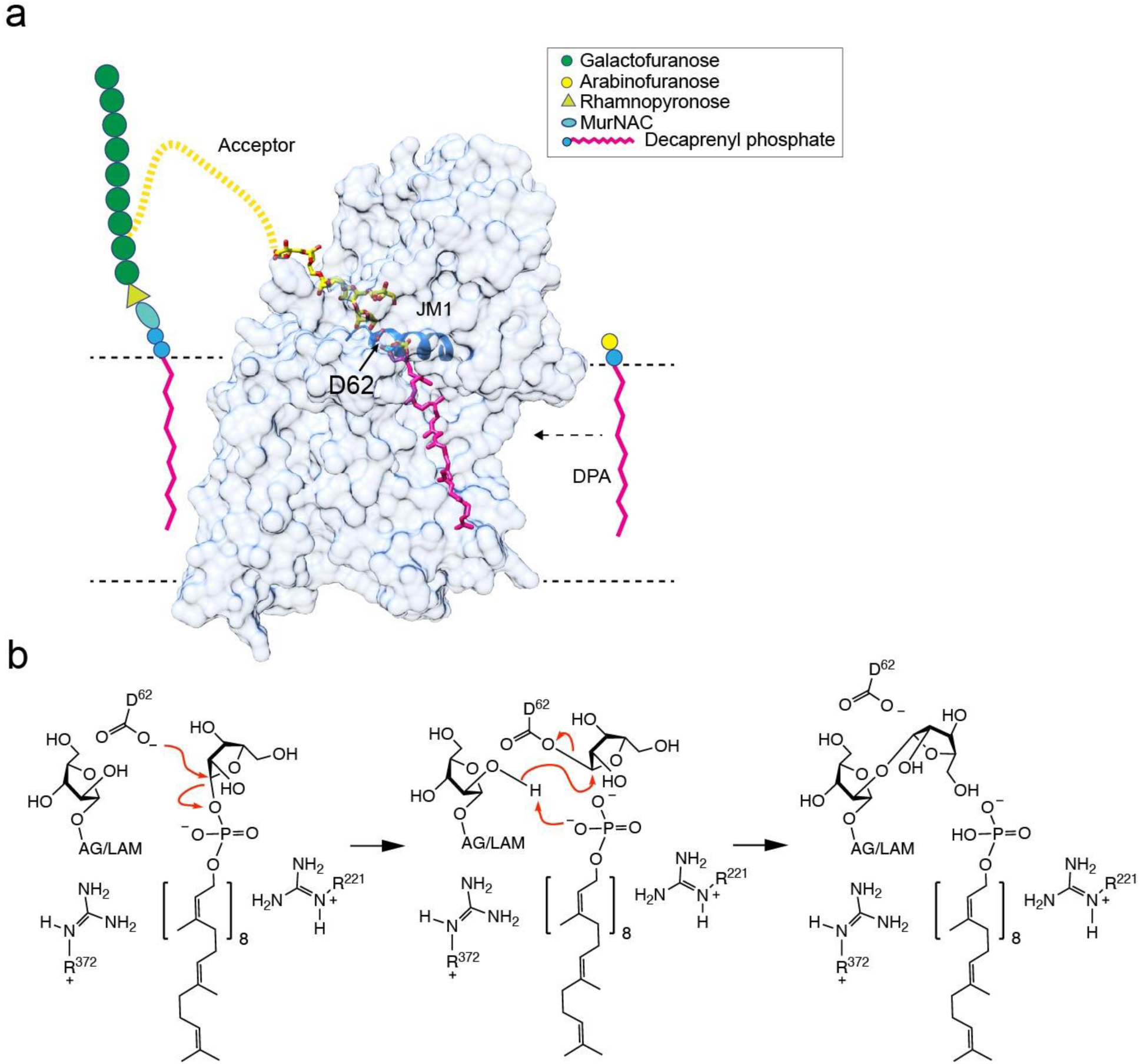
Proposed mechanism for AftB-catalyzed retaining arabinosyl transfer. (a). Schematic representation of AftB’s role in the biosynthesis of arabinogalactan. AftB directs the addition of arabinofuranosyl units from the donor molecule DPA onto the growing arabinan chain, maintaining the glycosidic bond stereochemistry. (b) Proposed catalytic model of AftB.

Given the retaining nature of AftB, we propose a double-displacement mechanism. Following our hypothesis, the catalytic cycle begins with Asp62 initiating a nucleophilic attack on the anomeric carbon of DPA (Fig. 5b), leading to the formation of a covalent β-Ara*f*–enzyme intermediate and inverting the stereochemistry of the Ara*f* residue to be transferred. Subsequently, the departing phosphate group of DPA deprotonates the C2 hydroxyl group of the acceptor substrate (Fig. 5b), enabling it to perform a nucleophilic attack on the anomeric carbon of the covalent β-Ara*f*–enzyme intermediate, breaking the arabinose–enzyme bond and forming a new β-(1→2) glycosidic bond. This second nucleophilic attack inverts the stereochemistry of the Ara*f* moiety once more, resulting in a net retention of the original β-configuration of the donor (Fig. 5b).

## DISCUSSSION

AG and LAM are complex polysaccharides that are major constituents of the cell envelope of mycobacteria. We have determined structures of AftB, an arabinofuranosyltransferase responsible for adding the terminal Ara*f* residues to both AG and LAM, in its apo and 2F-FPA bound state using cryo-EM to a resolution of 2.9 Å and 3.4 Å, respectively. These structures reveal that AftB belongs to the GT-C_A_ subclass of GT-C glycosyltransferases, with a conserved structural core of 7 TM helices. They also show an irregular tube-shaped cavity that bridges the two proposed substrate binding sites. Based on the structural characteristics of the cavity and our experimental evidence, we can propose how the two substrates bind, and a mechanism of action.

The donor substrate, DPA, is a lipid-linked Ara*f* that is anchored to the membrane via a decaprenyl-phosphate moiety. Based on the characteristics of the donor substrate and the biophysical properties of the cavity observed in the structure, we propose that the lipid tail of DPA extends out of the cavity towards TM helices 6 and 7 and into the lipid bilayer, allowing its reactive Ara*f*– phosphate moiety to be oriented towards the catalytic core. The acceptor substrates for AftB are the elongated AG and LAM polysaccharides, which are both connected to the membrane via a galactan or mannan core, respectively. Both AG and LAM share common motifs at the non-reducing end of their arabinan domains that serves as the acceptor site for AftB. The irregular shape of the catalytic cavity and its wider periplasmic-facing opening appear to be adapted to accommodate the arabinan domains of AG and LAM to access the active site from the periplasm. While AG and LAM share a similar branched motif at the non-reducing end of their arabinan domains, LAM possesses a more complex arabinan structure that also includes linear arabinan regions. The architectural features of the catalytic cavity of AftB appear well-suited for accommodating both linear and branched terminal arabinan acceptor motifs shared by AG and LAM.

Glycosyltransferases can be classified as either inverting or retaining based on the stereochemical outcome of the glycosyl transfer reaction^50,59^. The former inverts the configuration of the anomeric carbon of the donor substrate in the resulting product^48,60^, while the latter maintains the configuration^48,60,61^. AftB is a retaining glycosyltransferase and, to the best of our knowledge, is the first structurally characterized retaining GT-C fold glycosyltransferase.

The catalytic mechanism of retaining glycosyltransferases is a subject of debate^62^, with two main mechanisms proposed: the front-face S_N_i (substitution nucleophilic internal) mechanism^61,63^ and the double-displacement mechanism^50,61,62^. Most structurally characterized retaining glycosyltransferases lack a suitably positioned catalytic base, supporting the front-face S_N_i mechanism^50,63,64^. However, recent studies of a retaining glycosyltransferase WbbB, which belongs to the GT-B fold and synthesizes 3-deoxy-D-manno-oct-2-ulosonic acid (Kdo) β-glycosides, provides evidence for a double-displacement mechanism^65–67^.

Our structural analysis of AftB in complex with the donor analog 2F-FPA reveals that the putative catalytic residue Asp62, which belongs to the conserved DD motif, is located near the anomeric carbon of the donor analog, poised to form a covalent intermediate. We attempted to obtain a ternary structure including both the donor analog and an acceptor substrate analog, which would have provided additional insights into the catalytic mechanism. However, despite our efforts, we could not successfully capture this entity. Despite the absence of a ternary structure, the presence of an appropriately placed catalytic base in the active site, as observed in our donor analog complex, suggests that AftB uses a double-displacement mechanism. Notably, the replacement of this aspartate residue with alanine abrogates function, implicating it in catalysis.

The mycobacterial cell envelope plays a crucial role in the resilience and virulence of the pathogen. Targeting the biosynthesis of two key components of this envelope, AG and LAM, has proven to be a successful strategy for anti-TB drug development, as exemplified by the front-line drug ethambutol, which inhibits the arabinosyltransferases EmbB and EmbC involved in arabinan domain assembly^21,68^. AftB catalyzes the final step in the biosynthesis of the arabinan domains in both AG and LAM and therefore represents a potential target for anti-TB drug development^36,40^. Notably, AftB is highly conserved across mycobacterial species, including non-tuberculous mycobacteria (NTM) such as *M*. *chubuense* (studied herein), *M*. *abscesses*, and *M*. *chelone*^69–71^ (Extended Data Fig. 8), which cause various respiratory and skin infections. The conservation suggests AftB as a potential target for broad-spectrum anti-TB agents against multiple mycobacterial pathogens. Our study should help lay the groundwork for the rational design of AftB inhibitors and thereby hold promise for the development of novel therapeutics, which are urgently needed in the face of escalating drug resistance in both TB and NTM infections^26,69^.

## METHODS

### Overexpression and purification of *Mycobacterium chubuense* AftB

Wild type *Mycobacterium chubuense* AftB was cloned into pNYCOMPSC23 plasmid^72^. The procedure for design and synthesis of pNYCOMPSC23 plasmid was described in detail in a previous protocol^72^. The plasmids were transformed into BL21 (DE3) pLysS *E. coli* for protein expression. The following day, a single colony was selected to inoculate a starter culture consisting of 50 mL Terrific Broth (TB) medium (Fisher), supplemented with 50 μg/mL ampicillin and 35 μg/mL chloramphenicol. The starter culture was grown overnight at 37°C with shaking at 220 rpm in an incubator shaker (New Brunswick Scientific). The next day, four flasks, each containing 1 L of TB medium with 50 μg/mL ampicillin and 35 μg/mL chloramphenicol, were inoculated with 10 mL of the starter culture and grown at 37°C with shaking at 220 rpm until the OD600 reached 0.6–0.8. The cultures were then cooled to 22°C, and protein expression was induced by adding 0.5 mM IPTG for 16 hours. Cells were harvested by centrifugation at 4,000 x g in H6000A/HBB6 rotor (Sorvall) for 30 min at 4°C, and the pellet was resuspended with lysis buffer containing 20 mM HEPES pH 7.5, 200 mM NaCl, 20 mM MgSO_4_, 10 mg/mL DNase I (Roche), 8 mg/mL RNase A (Roche), 1 mM tris(2-carboxyethyl) phosphine hydrochloride (TCEP), 1 mM PMSF, 1 tablet/1.5 L buffer EDTA-free cOmplete protease inhibitor cocktail (Roche). Cells in lysis buffer were lysed by passing through a chilled Emulsiflex C3 homogenizer (Avestin) three times. The lysate was centrifuged at 3,000 x g in a Centrifuge 5810 R (Eppendorf) at 4°C for 5 min to remove cell debris and non-lysed cells. To isolate the cell membrane, the supernatant was ultracentrifuged in a Type 45 Ti Rotor (Beckman Coulter) at 185,600 x g for one hour. For 4 L of culture, the yield was approximately 4 g of membrane pellet, which was resuspended in the lysis buffer up to 80 mL and homogenized using a handheld glass homogenizer (Kontes) on ice. The membrane fraction was stored at −80 °C until further use. The membrane fraction was thawed and solubilized by adding n-dodecyl-b-D-maltopyranoside (DDM) to a final concentration of 1% (w/v) and was rotated gently at 4 °C for 2 hours. Insoluble material was removed by ultracentrifugation at 185,600 x g in a Type 45 Ti Rotor at 4 °C for 40 min. The supernatant was added to preequilibrated Ni^2+^-NTA resin (QIAGEN) in the presence of 40 mM imidazole and incubated with gentle rotation at 4 °C for 2 hours.

The resin was then washed with 10 column volumes of wash buffer containing 20mM HEPES pH 7.5, 200 mM NaCl, 65 mM imidazole, 0.1% DDM and eluted with 3 column volumes of elution buffer containing 20 mM HEPES pH 7.5, 200mM NaCl, 300mM imidazole, 0.03% DDM. The eluted protein was exchanged into a buffer containing 20 mM HEPES pH 7.5, 200mM NaCl, 0.03% DDM using a PD-10 desalting column (GE).

### Preparation of Mycobacterial Membranes for Arabinosyltransferase Assay

Wild-type *M. smegmatis* mc^2^155 cells were grown in 50 mL of Luria–Bertani (LB) broth for 72 h at 37 °C and 200 rpm. An Erlenmeyer flask with 2 L of LB broth was inoculated with 10 mL of the saturated starter culture and incubated for 48 h at 37 °C and 130 rpm. Cells were harvested by centrifugation at 2,600 x g for 15 min at 4 °C, resuspended in ice-cold buffer A [50 mM MOPS (pH 7.9), 5 mM 2-mercaptoethanol and 10 mM MgCl_2_], and disrupted by passing the suspension through a high-pressure homogenizer at 20,000 psi. The lysate was centrifuged at 27,000 x g for 30 min at 4 °C. The cell wall pellet was removed, and the supernatant was recentrifuged at 100,000 x g for 2 hours at 4 °C. The supernatant was discarded, and the pellet of enzymatically active membrane was gently resuspended in 1 mL of buffer A with gentle homogenization using a Dounce homogenizer. The protein concentration of the membrane fractions was determined using the Pierce BCA protein assay kit and was typically between 15 and 20 mg/mL.

### Arabinosyltransferase Assay

The arabinosyltransferase activity was assessed using wild-type *M. smegmatis* membranes and *E. coli* membranes expressing *Mc*AftB as described above. Untransformed *E. coli* membranes were used as a negative control. A typical reaction mixture contained FPA (1 mM), synthetic acceptor (0.2 mM), ATP (1 mM), DMSO (2% v/v), buffer A [50 mM MOPS (pH 7.9), 5 mM 2-mercaptoethanol and 10 mM MgCl_2_] and the respective membranes (2 mg) in a total volume of 200 μL. The reaction mixture was incubated at 37 °C overnight and then terminated by adding 200 μL of ethanol. The resulting mixture was centrifuged at 20,000 x g for 15 min, and the supernatant was loaded onto a pre-equilibrated strong anion exchanger (SAX) cartridge. The cartridge was eluted with 5 mL of 50% ethanol–water. The eluate was evaporated to dryness, and the dried enzymatic products were treated with Ac_2_O (100 µL) in pyridine (100 µL). The reaction mixture was stirred overnight at room temperature before being concentrated and then partitioned between the two phases (1:1) of chloroform and water. The organic layer was concentrated before being analyzed by MALDI mass spectrometry.

### Nanodisc reconstitution of AftB

The protein was incorporated into lipid nanodiscs with a molar ratio 1:5:250 of membrane scaffold protein 1E3D1 (MSP1E3D1): (POPG) 1-palmitoyl-2-oleoyl-sn-glycero-3-phospho-(1’-rac-glycerol) (Avanti), and incubated for 2 hours with gentle agitation at 4 °C. For the substrate bound structure, 2F-FPA was added to the reconstitution mixture at a molar ratio of 1:3 (AftB: 2F-FPA). The POPG lipid was prepared by adding the solid extract to deionized water to a final concentration of 20 mM. The mix was put on ice and then gently sonicated with a tip sonicator (Fisher Scientific) to dissolve the lipids until the mixture became semitransparent.

To initiate nanodisc reconstitution, 100 mg Biobeads (Bio-Rad) per mL of protein solution was added to the mixture and incubated by gentle rotation at 4 °C overnight. Biobeads were removed the next day by filtering the reconstitution mixture through an Ultrafree centrifugal filter unit (Fisher) at 16,100 x g in a Centrifuge 5415 R (Eppendorf) at 4°C for 1 min. The reconstitution mixture was rebound to Ni^2+^-NTA resin in the presence of 25 mM imidazole for 2 hours at 4 °C in order to remove free nanodisc. The resin was washed with 10 column volumes of wash buffer containing 20 mM HEPES pH 7.5, 200 mM NaCl and 50 mM imidazole, followed by 4 column volumes of elution buffer containing 20 mM HEPES pH 7.5, 200 mM NaCl and 300 mM Imidazole. The eluted protein was subsequently purified by size-exclusion chromatography (SEC) on a Superdex 200 Increase 10/300 GL column in buffer containing 20 mM HEPES pH 7.5, 200 mM NaCl.

### CryoEM sample preparation

Fractions containing AftB incorporated into nanodiscs were pooled and incubated with the Fab at 4 °C for 2 hours in a 1:3 molar ratio of protein to Fab. The protein mixture was further purified using a Superdex 200 Increase 10/300 GL SEC column in SEC buffer containing 20 mM HEPES pH 7.5, 200 mM NaCl.

For the apo structure, peak fractions were pooled and concentrated to 7 mg/mL using a 50 kDa cutoff filter concentrator (Amicon). The sample was frozen using a Vitrobot (Thermo Fisher) by applying 3 μL to a plasma cleaned (Gatan Solarus) 0.6/1-mm holey gold grid (Quantifoil UltrAuFoil). After a 30 s incubation, the grids were blotted using 595 filter paper (Ted Pella, Inc) for 8 s before being immediately plunged into liquid ethane for vitrification. The plunger was operated at 4 °C with greater than 90% humidity to minimize evaporation and sample degradation. For the substrate-bound structure, peak fractions were pooled and concentrated to concentration of 5 mg/mL. The sample was incubated with 2F-FPA at a 1:2 molar ratio of protein to substrate for one hour before being applied onto the grids. The grids were prepared in the same way as for the apo sample, but with a blotting time of 6 s.

### Data Collection

For the apo structure, images were recorded at the Columbia University Cryo-Electron Microscopy Center on a Titan Krios electron microscope (FEI), equipped with an energy filter and a K3 direct electron detection filter camera (Gatan K3-BioQuantum) using a 0.87 Å pixel size. An energy filter slit width of 20 eV was used during the collection and was aligned automatically every hour using the Leginon software package^73^. Data collection was performed using a dose of around 58 e^−^ per Å^2^ across 50 frames (50 ms per frame) at a dose rate of approximately 16 e^−^ per pixel per second, using a set defocus range of −1.2 μm to −2.2 μm. A total of 7,164 micrographs were collected over a single two-day session.

For the substrate-bound structure, images were recorded at the New York Structural Biology Center (NYSBC) on a Titan Krios (FEI; NYSBC Krios 2) operating at 300 kV equipped with a spherical aberration corrector, an energy filter (Gatan GIF Quantum), and a post-GIF K2 Summit direct electron detector, using a 0.8460 Å pixel size. An energy filter slit width of 20 eV was used during the collection and was aligned automatically every hour using the Leginon software package^73^. Data collection was performed using a dose of around 50.3 e^−^ per Å^2^ across 24 frames (50 ms per frame) at a dose rate of approximately 41 e^−^ per pixel per second, using a set defocus range of −0.8 μm to −2.5 μm. A total of 16,290 micrographs were collected over a single session of three days and 20 hours (92 hours total).

### Identification of *Mc*AftB-specific Fab using phage display

AftB was reconstituted into chemically biotinylated MSP1E3D1 as previously described^44,74^. Selection for Fabs was performed starting with Fab Library E^45,75^. Targets and the library were first diluted in selection buffer (20 mM HEPES, pH 7.4, 150 mM NaCl and 1% BSA). Five rounds of sorting were performed using a protocol adapted from published protocols^76,77^. In the first round, bio-panning was performed manually using 400 nM of AftB, which was first immobilized onto magnetic beads and washed three times with selection buffer. The library was incubated for one hour with the immobilized target, beads were subsequently washed three times with selection buffer, and then beads were used to directly infect log-phase *E. coli* XL-1 Blue cells. Phage were amplified overnight in 2XYT media supplemented with ampicillin (100 µg/mL) and M13-K07 helper phage (10^9^ pfu/mL). To increase the stringency of selection pressure, four additional rounds of sorting were performed by stepwise reduction of the target concentration: 200 nM in the 2^nd^ round, 100 nM in the 3^rd^ round, and 50 nM in the 4^th^ and 5^th^ rounds. These rounds were performed semi-automatically using a KingFisher magnetic beads handler (Thermo Fisher Scientific). For each round, the amplified phage population from each preceding round was used as the input pool. Additionally, amplified phage were precleared prior to each round using 100 µL of streptavidin paramagnetic particles, and 2.0 µM of empty MSP1E3D1 nanodiscs were used throughout the selection as competitors in solution. For rounds 2-5, prior to infection of log-phage cells, bound phage particles were eluted from streptavidin beads by 15 min incubation with 1% Fos-choline-12 (Anatrace).

### Single-point phage ELISA to validate Fab binding to *AftB*

96-well plates (Nunc) were coated with 2 µg/mL Neutravidin and blocked with selection buffer. Colonies of *E. coli* XL-1 Blue cells harbouring phagemids from the 4^th^ and 5^th^ rounds were used to inoculate 400 µL 2XYT media supplemented with 100 µg/mL ampicillin and 10^9^ pfu/mL M13-KO7 helper phage, and phage were subsequently amplified overnight in 96-well deep blocks with shaking at 280 rpm. Amplifications were cleared of cells with a centrifuge step and then diluted 10-fold into ELISA (selection buffer with 2% BSA). All phage were tested against wells with immobilized biotinylated MSP1E3D1-reconstituted AftB (30 nM), empty biotinylated-MSP1E3D1 nanodiscs (50 nM), or buffer alone to determine specific target binding. Phage ELISA was subsequently performed as previously described^74,76^ where the amount of bound phage was detected by colorimetric assay using an anti-M13 HRP-conjugated monoclonal antibody (GE Healthcare). Binders with high target and low non-specific signal were chosen for subsequent experiments.

### Fab cloning, expression and purification

Specific binders based on phage ELISA results were sequenced at the University of Chicago Comprehensive Cancer Center DNA Sequencing facility and unique clones were then sub-cloned into the Fab expression vector RH2.2 (kind gift of S. Sidhu) using the In-Fusion Cloning kit (Takara). Successful cloning was verified by DNA sequencing. Fabs were then expressed and purified as previously described^74^. Following purification, Fab samples were verified for purity by SDS-PAGE and subsequently dialyzed overnight in 20 mM HEPES, pH 7.4, 150 mM NaCl.

### Assessment of Fab binding affinity to AftB

To measure the apparent binding affinity, multi-point ELISAs using each purified Fab were performed in triplicate. Briefly, AftB (30 nM) or empty biotinylated-MSP1E3D1 nanodiscs (50 nM) were immobilized onto 96-well plates coated with Neutravidin (2 µg/mL). Fabs were diluted serially 3-fold into ELISA buffer using a starting concentration of 3 µM, and each dilution series was tested for binding to wells containing either AftB, empty nanodiscs, or no target at all. The Fab ELISA was subsequently performed as previously described^76^, where the amount of bound Fab was measured by a colorimetric assay using an HRP-conjugated anti-Fab monoclonal antibody (Jackson ImmunoResearch). Measured A_450_ values were plotted against the log Fab concentration, and EC_50_ values were determined in GraphPad Prism version 8.4.3 using a variable slope model assuming a sigmoidal dose response.

### Single-particle cryo-EM data processing and map refinement

All data sets were corrected for beam-induced motion with Patch Motion Correction implemented in cryoSPARC v.2.15^78^ and the contrast transfer function (CTF) was estimated with Patch CTF. For the apo structure, 5.03 million particles were automatically picked using a blob-picker job and subjected to multiple rounds of 2D classification. Representative 2D classes of 118,434 particles clearly showing the Fab bound complex in the side views were selected as input for Topaz training. The resulting model was used to pick particles using Topaz Extract. Initially, 1.8 million particles were extracted in a box of 600 pixels and Fourier cropped to 150 pixels for initial cleanup. Initially, 2,790,710 particles were extracted with a box size of 320 pixels and binned four times. Multiple rounds of 2D classification were performed to clean up the particles using a batchsize per class of 400 and ‘‘Force Max over poses/shifts’’ turned off with 40 online-EM iterations. Classes of total 376,801 particles which display clear features of a Fab-bound nanodisc-embedded membrane protein were re-extracted using a 360-pixel box size without binning. *Ab initio* reconstruction was performed in cryoSPARC v.2.15 using two classes and a class similarity parameter of 0.1. One good class comprised of 241,095 particles was subjected to heterogeneous refinement and a final class comprising 192,312 particles was subjected to Nonuniform refinement and yielded a reconstruction with a resolution of 3.0 Å (FSC = 0.143). A subsequent local refinement was performed using a mask covering AftB and the variable region of the Fab and resulted in a density map at 2.85 Å resolution.

For the substrate bound structure, 5.29 million particles were automatically picked using a blob-picker job and subjected to multiple rounds of 2D classification. Representative 2D classes of 10,186 particles clearly showing the Fab bound complex in the side views were selected as input for Topaz training. The resulting model was used to pick particles using Topaz Extract. Initially, 2.01 million particles were extracted in a box of 320 pixels and Fourier cropped to 80 pixels for initial cleanup. Multiple rounds of 2D classification were performed to clean up the particles using a batchsize per class of 400 and ‘‘Force Max over poses/shifts’’ turned off with 40 online-EM iterations. Classes of total 279,138 particles which display clear features of a Fab-bound nanodisc-embedded membrane protein were reextracted using a 360-pixel box size without binning. *Ab initio* reconstruction was performed in cryoSPARC v.2.15 using two classes and a class similarity parameter of 0.1. One good class comprised of 183,657 particles was subjected to heterogeneous refinement and a final class comprised of 146,894 particles was subjected to Nonuniform refinement and yielded a reconstruction with a resolution of 3.8 Å (FSC = 0.143). A subsequent local refinement was performed using a mask covering AftB and the variable region of the Fab and resulted in a density map at 3.40 Å resolution.

### Model building

All model building was performed in Coot^79^. The apo AftB model was constructed *de novo* from the globally sharpened 2.85 Å map using the Phenix Map to Model tool^80^. Manual segment joining and residue assignment were guided by Xtalpred’s secondary structure prediction^81^. Ramachandran outliers were fixed manually in Coot before the structure was refined with Phenix real space^80^ refine with secondary structure and Ramachandran restraints

For substrate-bound AftB, the ligand files were generated in Coot, and similarly to the apo structure, manual model building, and refinement were conducted iteratively with Coot^79^ and Phenix^80^. The structural quality of both apo and substrate-bound forms was assessed using Molprobity^82^. For most of the Fab region, with the exception of the binding interface, modeling was based on an existing high-resolution structure (PDB ID 5UCB).

### Molecular dynamics simulations

The CG parameters for DPA and terminal-Ara*f*_4_ were generated based on previously published lipid^83^ and Martini 3 parameters^84–86^. The values for bond lengths and angles were generated from AT simulation data, where the molecules of interest were parameterized using CGenFF^87^, and PyCGTOOL^88^ was used to get exact values.

For CG simulations, the apo structure was converted to the Martini 3 forcefield using martinize2^89^ including a 1000 kJ mol^−1^ nm^−2^ elastic network. The structure was then embedded in a PE:PG (80:20) membrane using the insane script^90^. Each simulation box was then solvated and neutralized with 150 M NaCl. For DPA simulations, 2% DPA was added to the upper leaflet of the membrane. For the simulations including the terminal-Ara*f*_4_, two copies of coarse-grained substrate were added to the solvent in a random orientation and positioning. Systems were energy minimized using the steepest descents method. Production simulations used a timestep of 20 fs, and five simulations of 5 µs were performed for both DPA and terminal-Ara*f*_4_ containing systems. The C-rescale barostat^91^ was set at 1 bar and the velocity-rescaling thermostat^92^ was used at 310 K. All simulations were performed using Gromacs 2021.4^93,94^.

Coordinates for the atomistic systems were generated from the end snapshots of the CG simulations using CG2AT2^95^. Coordinates for both the substrates and products were built using the densities from the ligand-bound cryo-EM structure. For the substrate state the bound DPA analogue was modified to DPA and energy minimized. RoseTTAFold All-atom^55^ was used to fold and dock both substrates and products for comparison. Atomistic molecular simulations were performed with the CHARMM36m forcefield^96^, with parameters for the terminal-Ara*f*_4_ and terminal-Ara*f*_6_ (substrates and products respectively) created using CHARMM-GUI^97^. Parameters for the polyprenyl phosphate and DPA were created using CHARMM-GUI and combined with previous parameters from our past studies^98,99^. Energy minimizations of the systems were performed using the steepest descents method. A timestep of 2 fs was used for production simulations, with three independent simulations of 500 ns for substrate, product and apo states of AftB. The C-rescale barostat^91^ was set at 1 bar and the velocity-rescaling thermostat^92^ was used at 310 K. All simulations were performed using Gromacs 2021.4^93,94^.

For analysis, PyLipID^100^ was used to measure interactions between the substrates and the protein. Plumed^101^ was used in conjunction with matplotlib^102^ to generate the density plots. Visualization of the duration was performed with Visual Molecular Dynamics (VMD)^103^.

## ACKNOWLEDGMENTS

This work was supported by R35GM132120 (to FM). PJS acknowledges R01AI174416 (PI: M. Stephen Trent)), Wellcome (208361/Z/17/Z), MRC, BBSRC (BB/P01948X/1), and the Howard Dalton Centre for funding. PJS and CMB acknowledge Sulis at HPC Midlands+, which was funded by the EPSRC on grant EP/T022108/1, and the University of Warwick Scientific Computing Research Technology Platform for computational access. CMB was supported by an MRC studentship (MR/N014294/1). TLL thanks Academia Sinica intramural funding and the Canadian Glycomics Network (SD-1) for support. This project made use of time on ARCHER2 and JADE2 granted via the UK High-End Computing Consortium for Biomolecular Simulation, HECBioSim (http://www.hecbiosim.ac.uk), supported by EPSRC (grant no. EP/R029407/1). MA acknowledges support by FCT -Fundação para a Ciência e a Tecnologia, I.P., through PTDC/BIA-BQM/4056/2020, MOSTMICRO-ITQB R&D Unit (DOI 10.54499/UIDB/04612/2020; DOI 10.54499/UIDP/04612/2020) and LS4FUTURE Associated Laboratory (DOI 10.54499/LA/P/0087/2020); and EU H2020 grant No 823780. Mass spectrometry data for products from the funtinal assay were acquired at the Medicinal Chemistry and Analytical Core Facilities in the Biomedical Translation Research Center located at National Biotechnology Research Park, supported by Academia Sinica Core Facility and Innovative Instrument (AS-NBRPCF-111-201). AAK acknowledges support from NIH grant R01GM117372.

## AUTHOR CONTRIBUTIONS

Y.L. and B.K. conducted the initial high-throughput screening and J.R. contributed to these early stages of the project. Y.L. performed molecular cloning, protein expression, purification, cryo-EM sample preparation, data processing, model building, and structural refinement. Y.C.S. and Y.J.W. conducted the AftB functional assay. P.S.T. and N.H.D. synthesized FPA for the functional assay. C.R.H. and H.Y.C. synthesized the Ara*f*_4_ acceptor for the functional assay. Contributions related to the functional assay were made under the guidance of T.L.L. C.M.B. and P.J.S. carried out all molecular dynamics simulations and docking studies. S.E., P.T., and A.A.K. identified, characterized, and purified the Fabs. Y.L. wrote the manuscript with critical input from R.N., F.M., P.J.S., C.M.B., M.A., T.L.L., and S.E. F.M., R.N. and P.J.S supervised the project. All authors reviewed and approved the final version of the manuscript.

## DATA AND MATERIALS AVAILABILITY

