## Supplementary material for "Structural insights into terminal arabinosylation biosynthesis of the mycobacterial cell wall arabinan"

### EXTENDED DATA FIGURE LEGENDS

Extended Data Figure 1

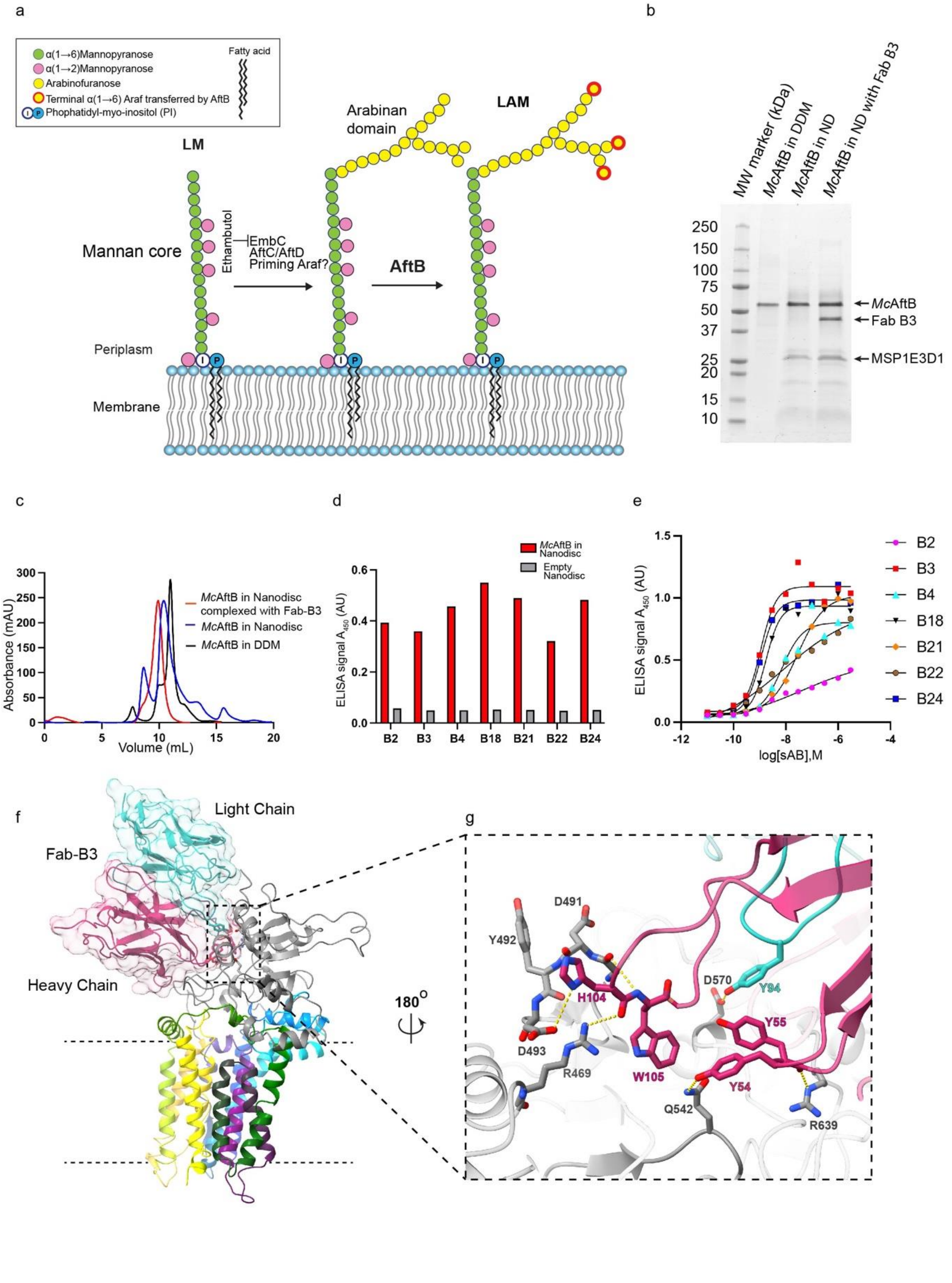

**Extended Data Fig. 1 Characterization and structural analysis of AftB.**

(a) Schematic representation of biosynthesis of arabinan domain of LAM catalyzed by AraTs, highlighting the role of the terminal arabinose addition catalyzed by AftB. (b) SDS-PAGE gel of AftB purification. The first lane shows AftB purified in DDM, the second lane shows AftB reconstituted into nanodiscs (using MSP1E3D1 and POPG), and the third lane shows AftB reconstituted into nanodiscs (MSP1E3D1 and POPG) with Fab-B3 bound. (c) SEC elution profile of purified AftB in detergent (black), incorporated into a nanodisc (blue), and incorporated into a nanodisc with Fab-B3 bound (red). (d) Single-point ELISA measuring the binding of phage-displayed sABs to AftB in MSP1E3D1 nanodiscs (red), empty nanodiscs (light grey). ELISA signal was measured at 450 nm absorbance. (e) Multi-point sAB ELISA: EC50 estimation for purified sAB binding to AftB incorporated into MSP1E3D1 nanodiscs, showing binding of B2, B3, B4, B18, B21, B22, B24. (f) AftB-Fab-B3 complex structure shown in ribbon with AftB's TM domain in rainbow colors and PD in grey. The Fab's light chain is depicted in teal, and the heavy chain in pink. (g) Magnified view of the interface between AftB and Fab-B3, highlighting the key interactions.

Extended Data Figure 2

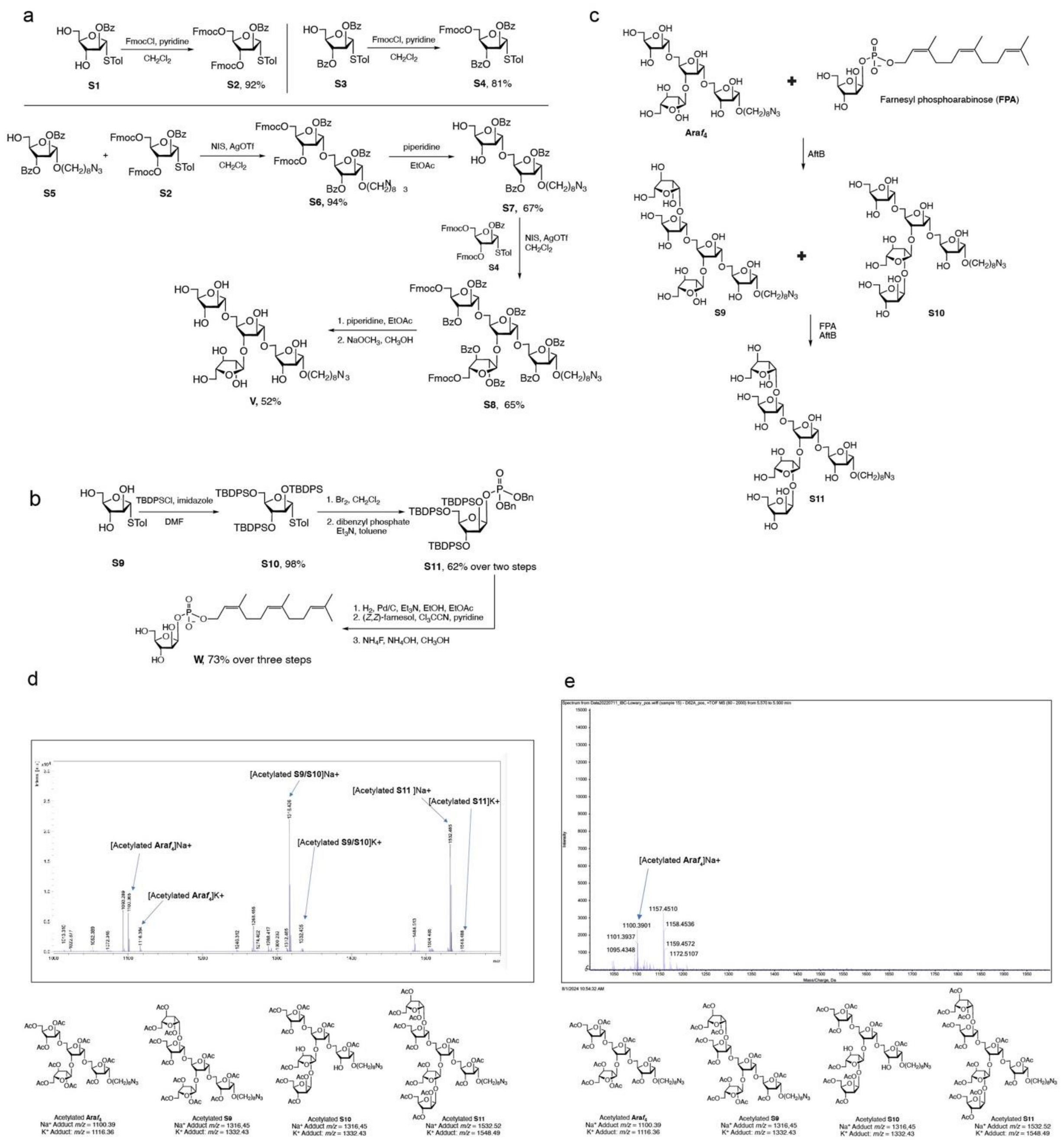

**Extended Data Fig. 2 Substrate synthesis and AftB enzymatic activity.**

(a) Synthesis of Ara*f*$_4$ and (b): Synthesis of FPA (c) AftB-catalyzed arabinofuranosylation of acceptor Ara*f*$_4$. The first formed products are pentasaccharide S9 and S10 and then hexasaccharide

S11. (d) Assay mixture produced by including acceptor Ara*f*$_4$, FPA, and *E. coli* membranes containing recombinant wild type AftB, followed by passage through an ion exchange cartridge and acetylation of the eluant. (e) Assay mixture produced by incubating acceptor Ara*f*$_4$, FPA, and *E. coli* membranes expressing recombinant AftB mutant D62A, followed by passage through an ion exchange cartridge and acetylation of the eluant. The absence of product formation in the D62A mutant demonstrates the critical role of Asp62 in AftB's catalytic activity.

Extended Data Figure 3

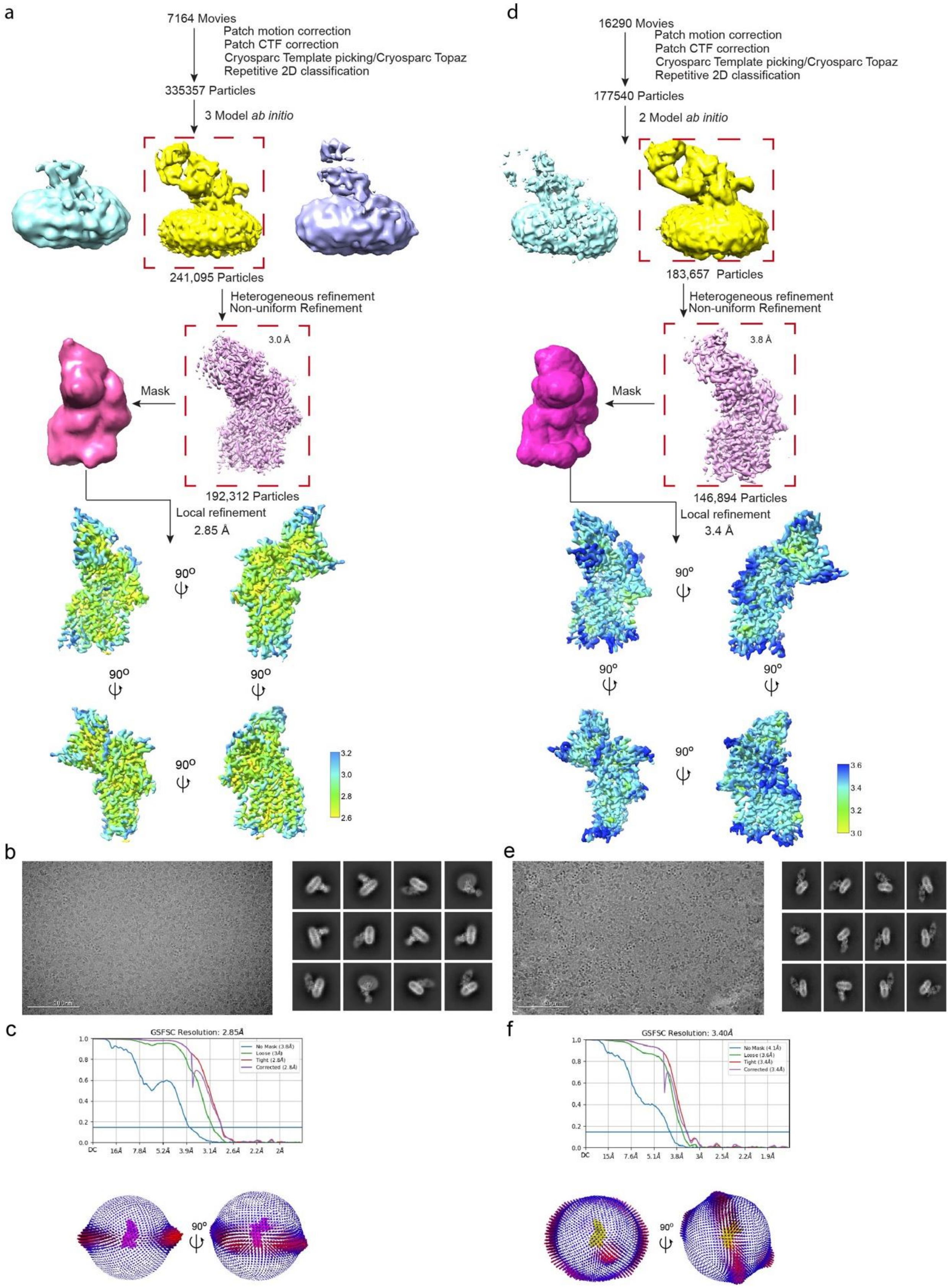

**Extended Data Fig. 3 Cryo-EM analysis of *Mc*AftB.**

(a) Schematic depiction of the cryo-EM data processing and structure determination for nanodisc-reconstituted apo *M.chubuense* AftB complexed with Fab-B3 using cryoSPARC. (b) Representative micrograph and representative 2D classes. (c) Top: Fourier shell correlation (FSC) curves of 3D reconstruction. Bottom: Euler angle distribution of particles used in the final 3D reconstruction. Final map shown in magenta. Each orientation is represented by a cylinder, with each cylinder's height and color (from blue to red) proportional to the number of particles for that specific direction. (d) Schematic depiction of the cryo-EM data processing and structure determination for 2F-FPA bound AftB complexed with Fab-B3 using cryoSPARC. (e) Representative micrograph and representative 2D class. (f) Top: Fourier shell correlation (FSC) curves of 3D reconstruction. Bottom: Euler angle distribution of particles used in the final 3D reconstruction. Final map is shown in magenta. Each orientation is represented by a cylinder, with each cylinder's height and color (from blue to red) proportional to the number of particles for that specific direction.

Extended Data Figure 4

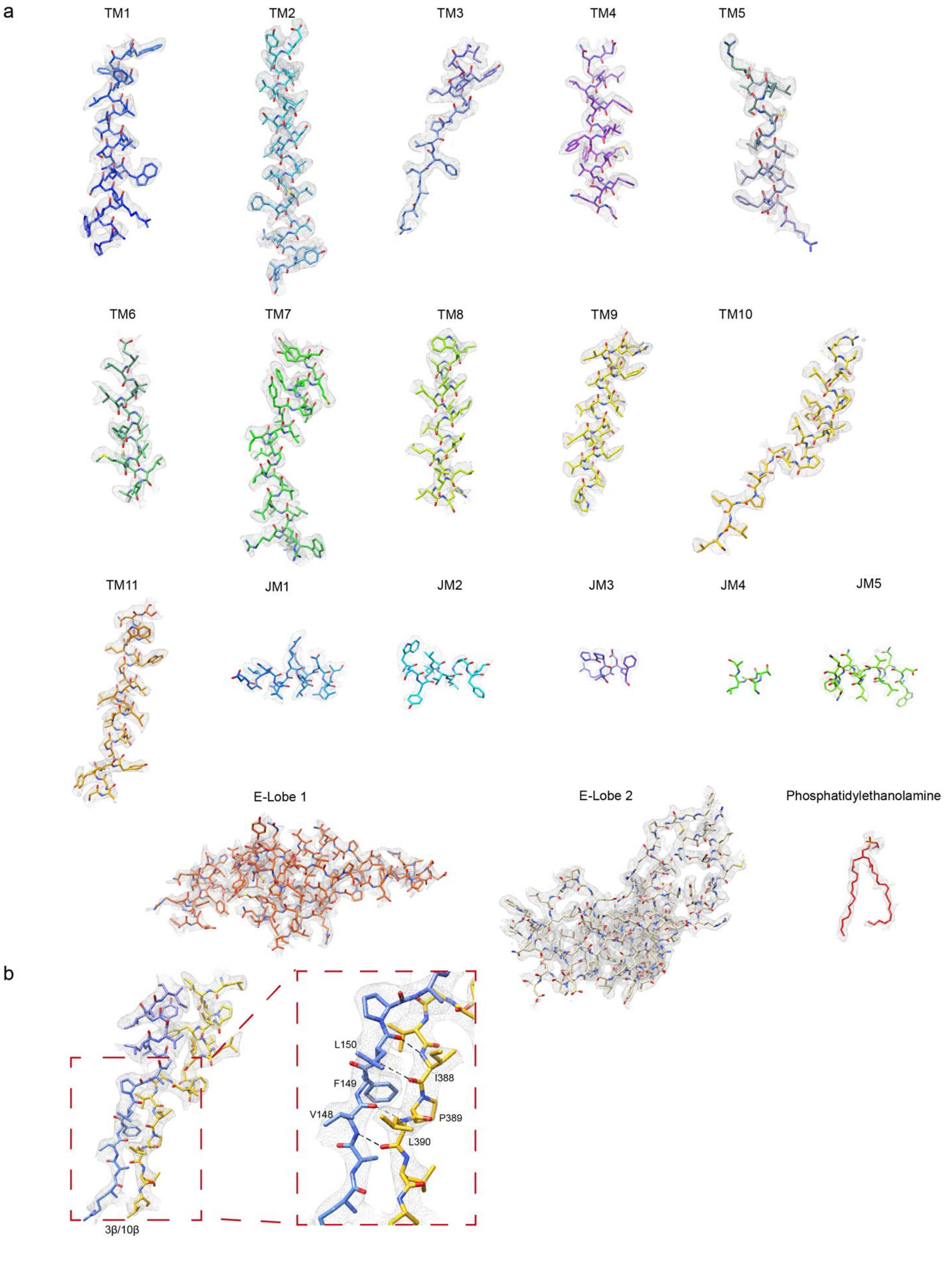

**Extended Data Fig. 4 EM Density of apo *Mc*AftB.**

(a) The atomic model of the structure of *Mc*AftB is colored in rainbow and rendered as a cartoon, with the side chains rendered as sticks. The map density is displayed as a mesh. (b) Density for 3β-10β, hydrogen bond is indicated with dashed lines.

Extended Data Figure 5

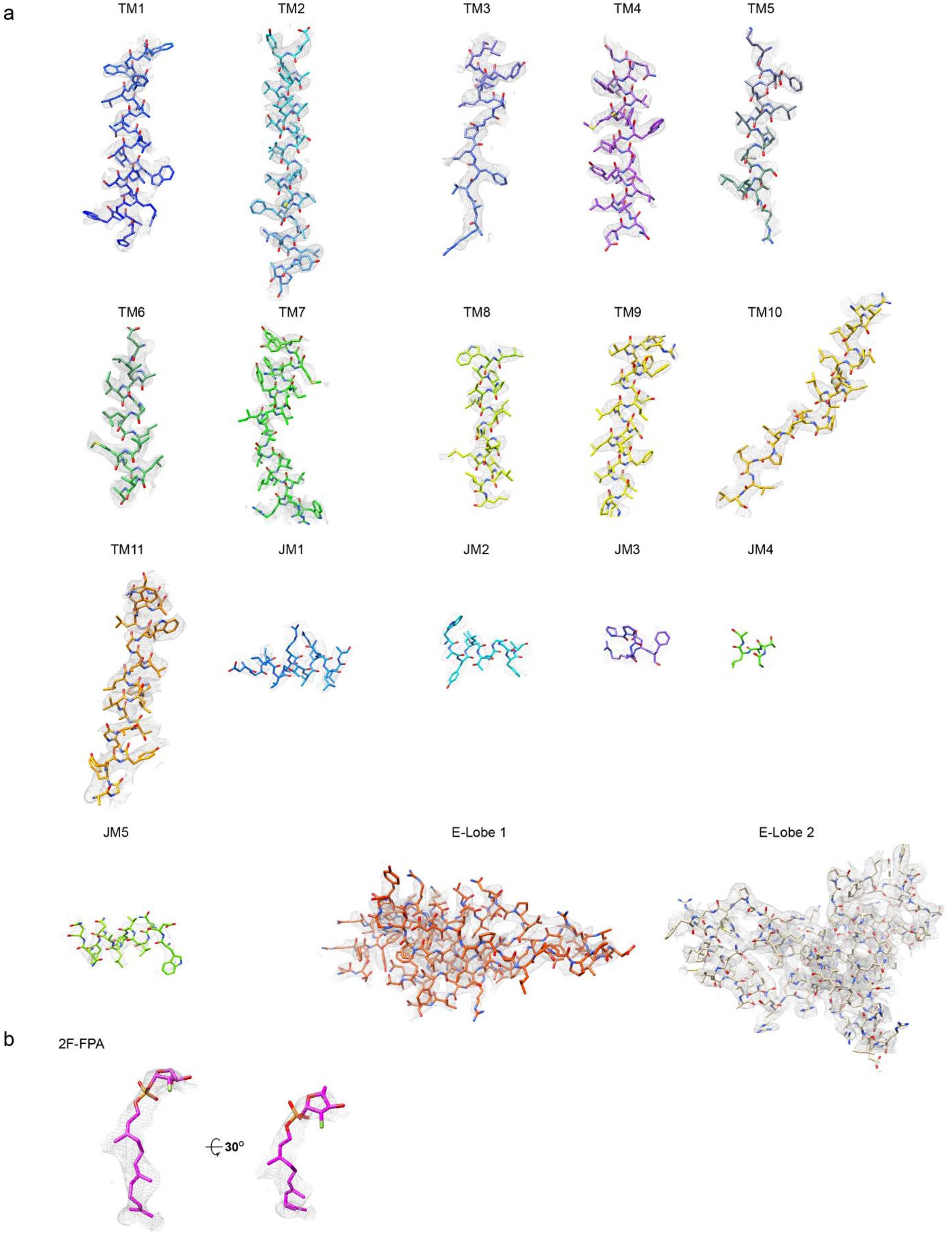

**Extended Data Fig. 5 EM Density of 2F-FPA bound *Mc*AftB.**

(a) The atomic model of the structure of AftB is colored in rainbow and rendered as a cartoon, with the side chains depicted as sticks. The map density is displayed as a mesh. (b) Density for 2F-FPA is displayed as mesh shown in two different views.

Extended Data Figure 6

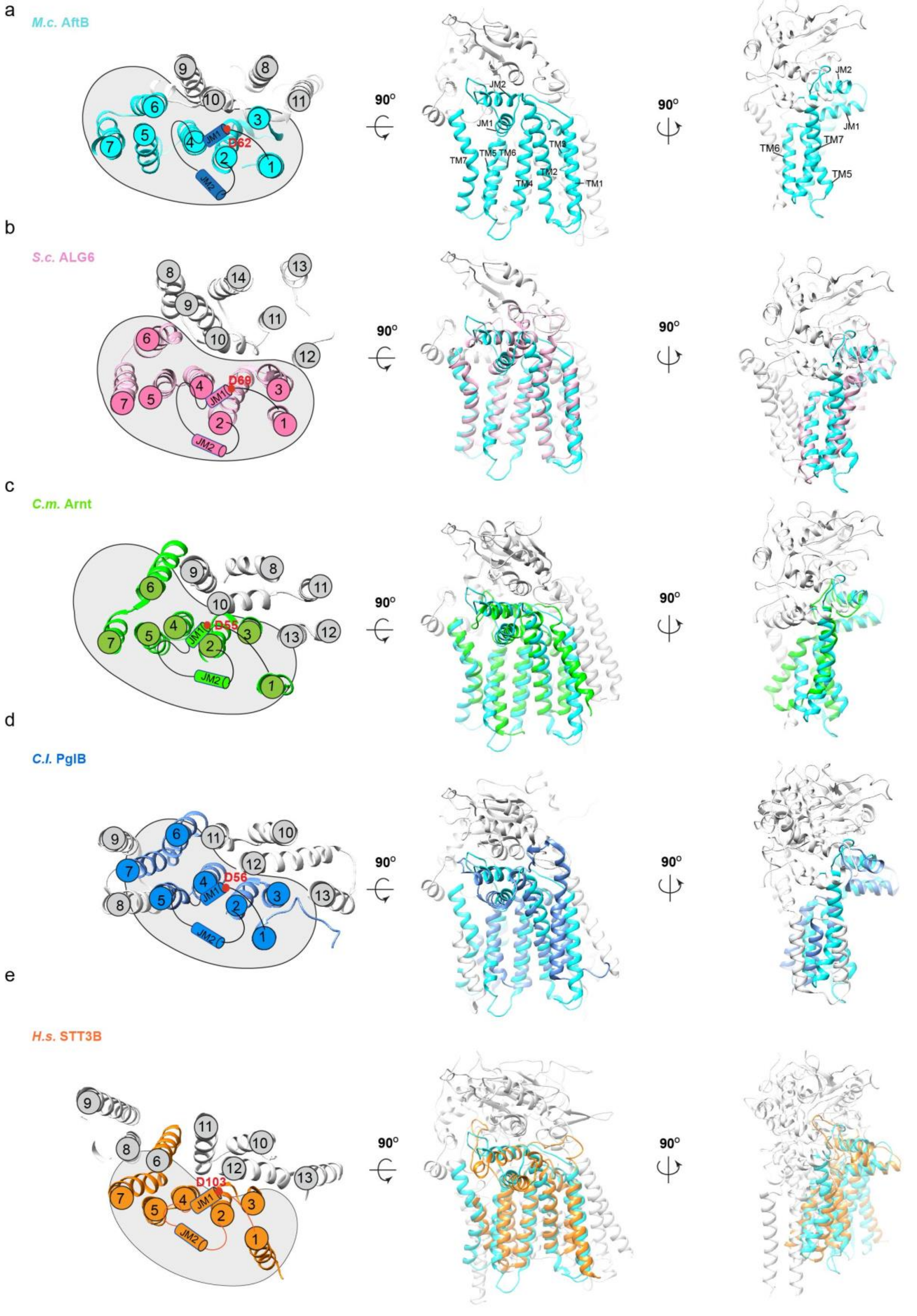

**Extended Data Fig. 6 Overall structure and comparison of *Mc*AftB with other GT-C fold glycosyltransferases.**

(a) Left, Top-down view of the TM domain of *Mc*AftB, highlighting the TM helices' spatial arrangement, with the first seven conserved helices colored in cyan, others in grey. TM helices are numbered with circles, JM1 and JM2 are represented as cylinders, and the catalytic residue is indicated by a red dot, in reference to Figure 1D. Middle and Right, two orthogonal views of *Mc*AftB. (b-e) Superimposition of *Mc*AftB with characteristic GT-C proteins from diverse species – *S. cerevisiae* ALG6 (PDB: 6SNH), *C. metallidurans* Arnt (PDB: 5F15), *C. lari* PglB (PDB: 5OGL), and *H. sapiens* STT3B (PDB: 6S7T) – illustrated in three distinct orientations, as presented in panel a.

Extended Data Figure 7

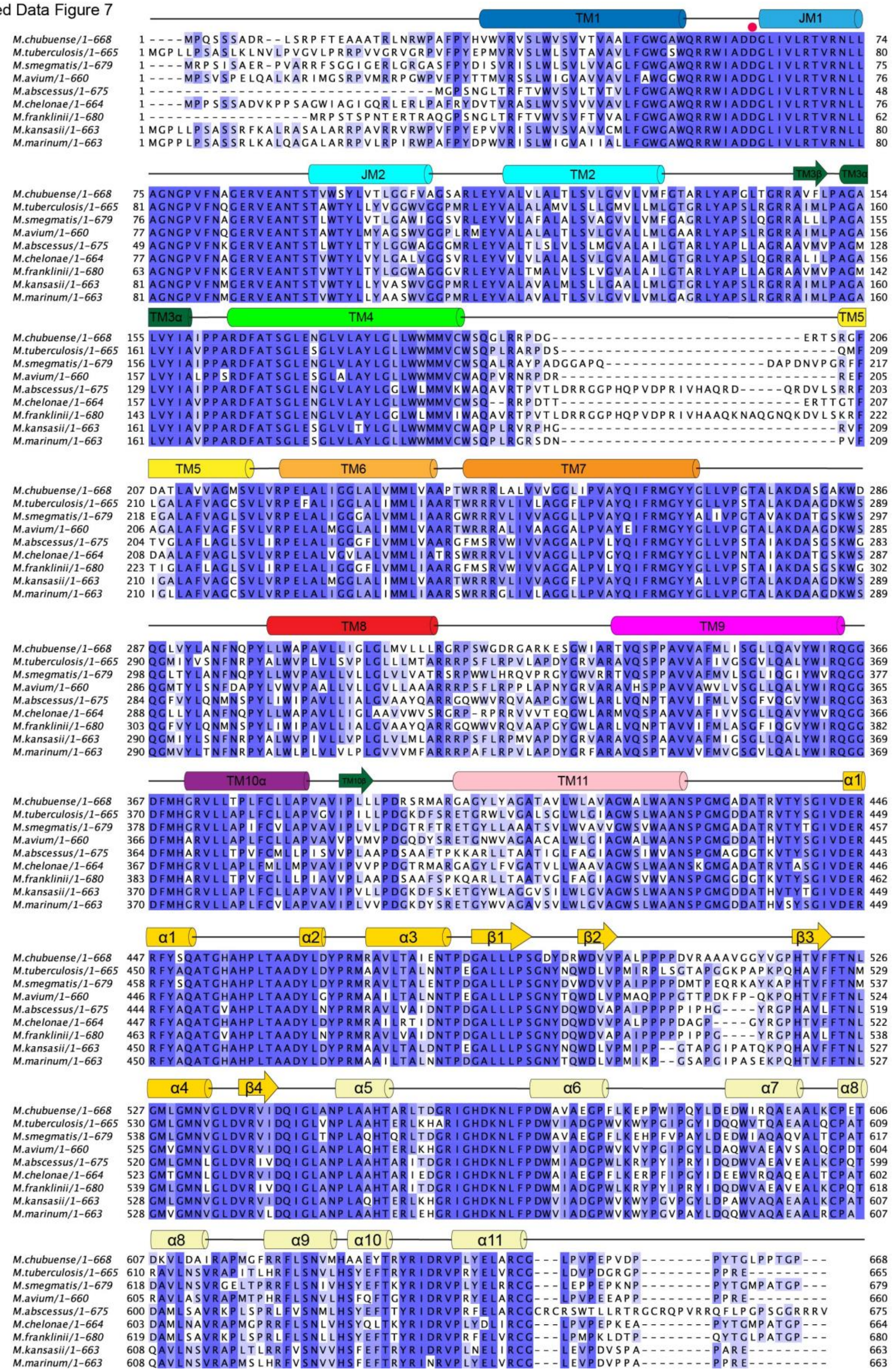

**Extended Data Fig. 7 Amino acid sequence conservation of AftB across mycobacterium species.**

Sequence alignment of AftB from *M. chubuense, M. tuberculosis, M. smegmatis, M. avium, M. abscessus, M. chelonae, M. franklinii, M. kansasii,* and *M. marinum*. The secondary structure elements, based on *M. chubuense* AftB, are indicated above the sequences. The structurally conserved and catalytically essential Asp62 residue is marked with a red dot.

Extended Data Figure 8

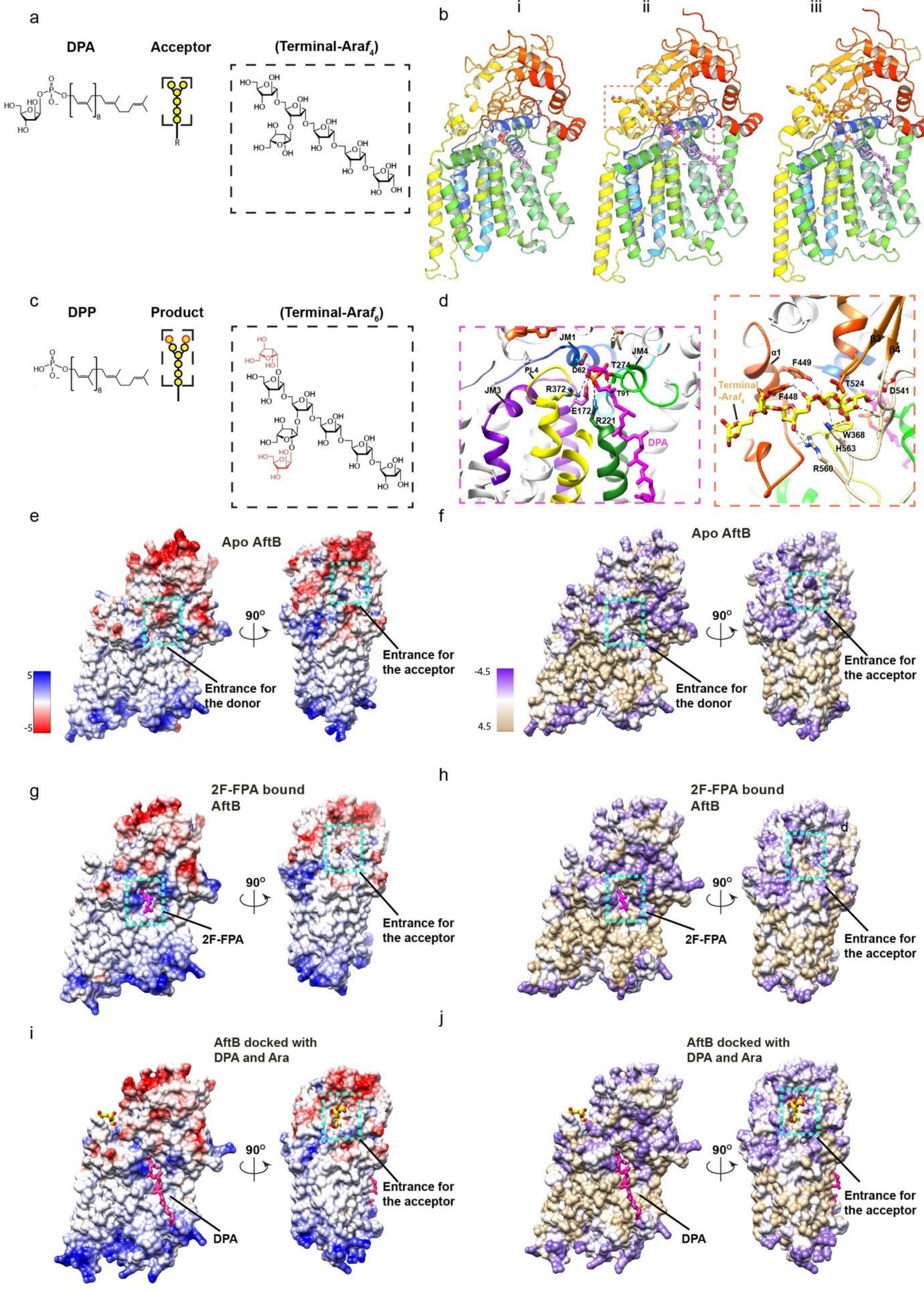

**Extended Data Fig. 8 Structural analysis and substrate binding models of AftB.**

(a) Donor substrate (DPA) and acceptor mimic (b) used in CG-MD simulations. (c) Computational models predicted by RoseTTAFold All-Atom docking[55]: (i) Structure of the apo *Mc*AftB solved by cryo-EM in this study. (ii) Computational model of AftB in complex with DPA and terminal-Ara*f*$_4$, predicted by RoseTTAFold docking. (iii) Computational model of AftB in complex with DP and terminal-Ara*f*$_6$, predicted by RoseTTAFold docking. (d) Detailed views of the substrate binding modes for the RoseTTAFold model shown in c (ii): Left: A magnified view of the binding mode for DPA in AftB showing key interacting residues. The catalytic residue Asp62 is positioned near the anomeric carbon of DPA. Coordinating residues for the phosphate group (Arg372, Arg221, Thr91, Thr243) are highlighted. Right: Magnified view of the proposed binding mode for the acceptor substrate in the PD of AftB. Key interacting residues are labeled. (e-f) Surface representation of apo *Mc*AftB colored by electrostatic potential (e) and hydrophobicity (f), shown in different orientations. Putative entrances for donor and acceptor substrates are highlighted with cyan dots. (g-h) Surface representation of 2F-FPA bound *Mc*AftB colored by electrostatic potential (g) and hydrophobicity (h), shown in different orientations. The 2F-FPA bound cavity and putative entrance for acceptor substrates are highlighted with cyan dots. (i-j) Surface representation of *Mc*AftB docked with DPA and simplified acceptor substrate terminal-Ara*f*$_4$, colored by electrostatic potential (i) and hydrophobicity (j), shown in different orientations. The cavities bound with DPA and terminal-Ara*f*$_4$ are highlighted with cyan dots. All the electrostatic potential was rendered on a range of ±5 kBT/e. All the hydrophobicity was colored using the kdHydrophobicity scale[106].

Extended Data Figure 9

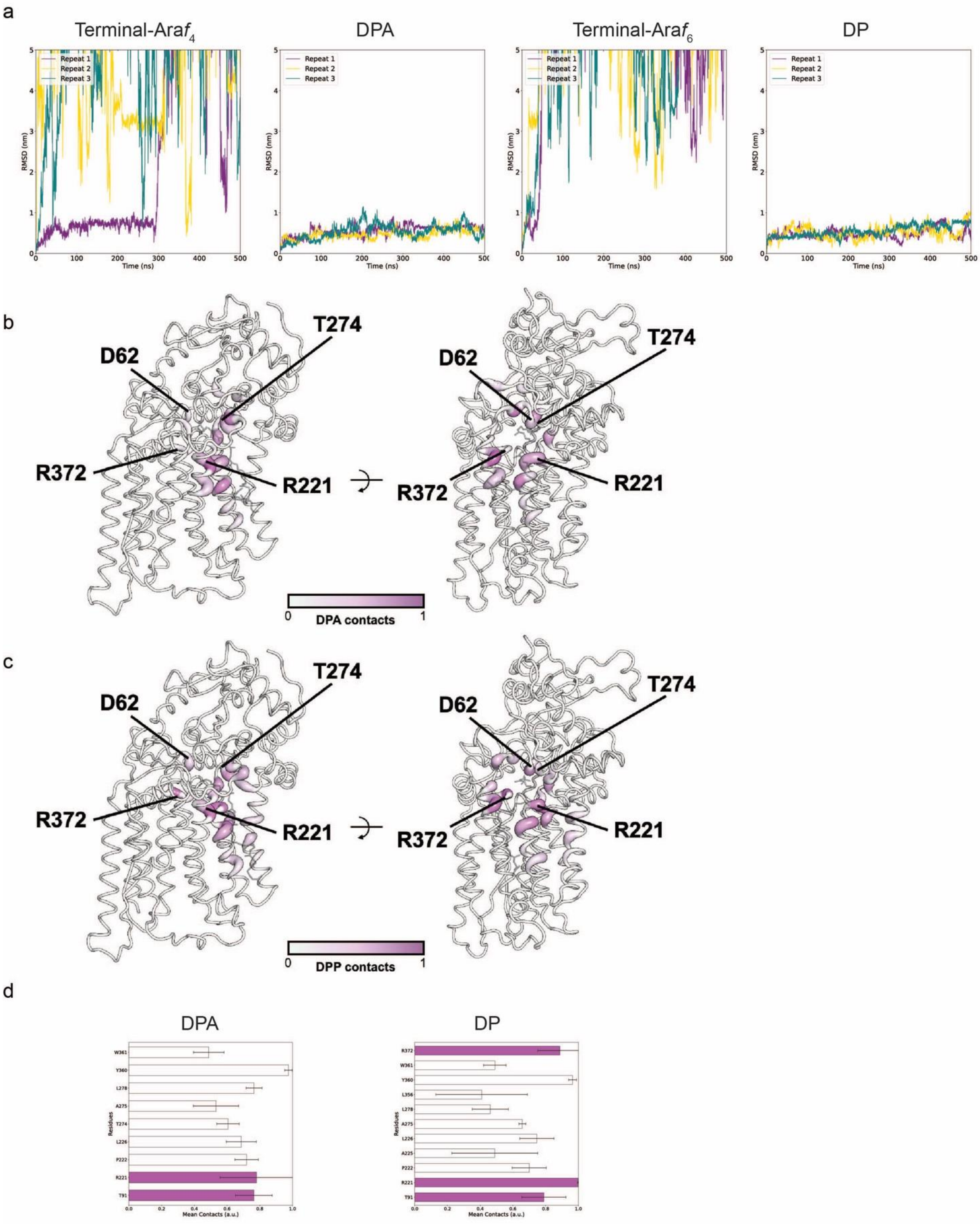

**Extended Data Fig. 9 Molecular dynamics analysis of AftB-substrate interactions.**

(a) The root mean squared deviation (RMSD) of DPA/DP and terminal-Ara$f_4$/ terminal-Ara$f_6$ with AftB. Each line represents an independent repeat. (b) Areas of AftB in contact with DPA in simulations. The darker the color, the more contacts throughout all simulations. The thickness of the cartoon also represents the number of contacts, where a contact value of 1 would represent contact with the ligand for the entire simulation. Key residue positions have been highlighted. (c) shows the same as (b), but for DP. (d) Contact graphs between AftB and DPA/DP. The mean contact value of three simulations is shown, with the error bar representing standard error. Residues that are in contact with 2F-FPA are highlighted. A contact value of 1 would represent contact with the ligand for the entire simulation, residues with contact values below 0.4 have been omitted for clarity.

Extended Data Figure 10

a

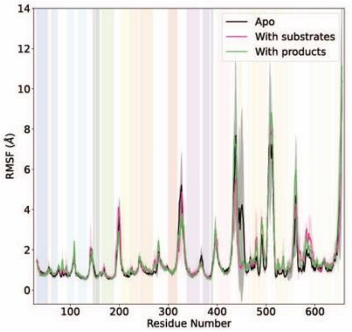


b

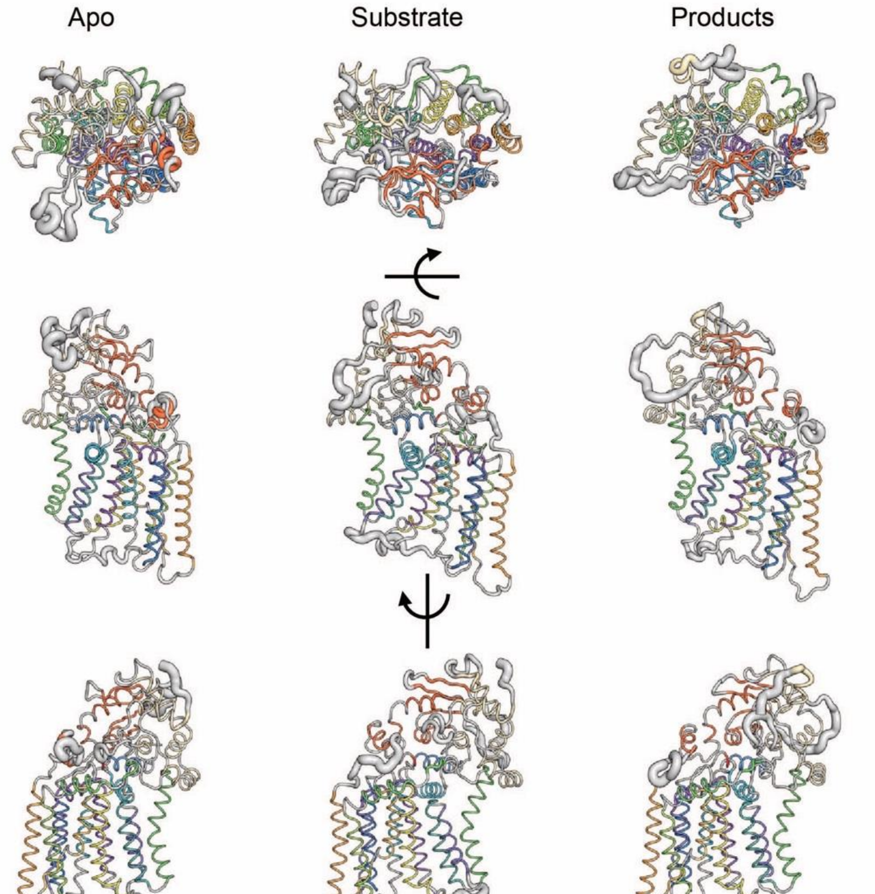


c

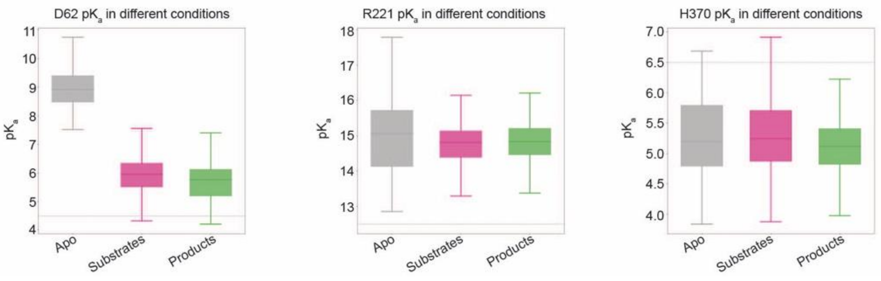

**Extended Data Fig. 10 Conformational Dynamics and p$K_a$ Analysis of AftB.**

The root mean squared fluctuation (RMSF) of AftB in the apo state, with substrates and with products. (a) A plot showing the RMSF over three simulations for each residue, with the outline signifying the standard error. The different topological regions of the protein as shown in Fig. 1c are highlighted by color. (b) The RMSF shown projected on the AftB structures, where the thicker regions are more dynamic in simulations. The helices are colored as in Fig. 1c. (c) Measured p$K_a$ values of selected residues in various simulation conditions. The expected value for that residue is shown as a gray line. The results shown include all independent repeats.

Table 1 . Cryo-EM data collection and modeling statistics

| | Apo AftB | AftB bound with 2F-FPA |
|---|---|---|
| Data Collection | | |
| Microscope | FEI Titan Krios-CEC | FEI Titan Krios-NYSBC |
| Camera | Gatan K3 | Gatan K3 |
| Voltage (kV) | 300 | 300 |
| Electron expose (e-/Å2) | 58 | 58 |
| Defocus range (μm) | -1.2 to -2.2 | -0.8 to -2.5 |
| Pixel size (Å) | 0.87 | 0.846 |
| Symmetry imposed | C1 | C1 |
| Initial particle images (No.) | 5,031,234 | 5,299,368 |
| Final particle images (No.) | 192,312 | 146,894 |
| Final Resolution (Å) | 2.85 | 3.4 |
| FSC threshold | 0.143 | 0.143 |
| Refinement | | |
| Model composition | | |
| Non-hydrogen atoms | 4741 | 4654 |
| Protein residues | 617 | 599 |
| Ligands | 0 | 1 |
| Waters | 0 | 0 |
| Mean B factor (Å2) | | |
| Protein | 63.5 | 61.2 |
| Ligands | | 45.9 |
| R.m.s. deviation | | |
| Bond lengths (Å) | 0.003 | 0.003 |
| Bond angles (°) Validation | 0.51 | 0.49 |
| Clashscore | 2 | 8 |
| Rotamers outlier (%) | 0 | 0 |
| Ramachandran plot | | |
| Favored (%) | 97 | 96.5 |
| Allowed (%) | 3 | 3.5 |
| Disallowed (%) | 0 | 0 |

#### Table

Cryo-EM data collection, refinement and validation statistics
